# PHATE-Derived Ancestry Coordinates Capture Continuous Population Structure in Human Genomic Data

**DOI:** 10.64898/2026.10.29.762525

**Authors:** Matthew Scicluna, Shuang Ni, César Miguel Valdez Córdova, Jean-Christophe Grenier, Raphaël Poujol, Mohamed Elyes Kanoun, João Felipe Rocha, Kevin R Moon, Sébastien Lemieux, Guy Wolf, Smita Krishnaswamy, Julie G Hussin

**Author notes:** These authors contributed equally to this work (Matthew Scicluna, Shuang Ni).

## Abstract

Population genetic data are increasingly visualized using low-dimensional embeddings as ancestry coordinates, yet how these methods represent population structure can depend strongly on sampling and construction choices. Principal component analysis (PCA), when visualized using only leading components, can obscure fine-scale structure, while nonlinear neighbor-based methods such as uniform manifold approximation and projection (UMAP) often fragment geographic and admixture gradients into discrete clusters. Here, we introduce a framework for constructing ancestry coordinates using potential of heat-diffusion for affinity-based trajectory embedding (PHATE), a diffusion-based method that preserves relationships across scales. Across synthetic benchmarks and large genomic datasets, we show that appropriate scale tuning and population balancing allow PHATE to recover geographic organization and admixture gradients. We show that incorrectly tuned neighborhood scale and population imbalance can distort the resulting representations of population structure. These results therefore provide a methodological framework and practical guidance for constructing ancestry visualizations that better reflect continuous human genetic diversity.

## 1 Introduction

Understanding the structure of human genetic variation can have broad implications for fields ranging from medicine to anthropology. Towards this goal, researchers have created population-genetic resources that are increasingly central to research efforts [1, 2]. Globally representative reference panels, such as the Human Genome Diversity Project (HGDP) [3] and the 1000 Genomes Project (1KGP) [4] intentionally sample populations across geographic regions to create catalogs of human genetic variation. More recently, large biobank cohorts, such as the UK Biobank (UKBB) [5] and All of Us (AoU) [6], have enabled population-genetic analyses at unprecedented scale within healthcare and national research settings.

To visualize the structure of these high-dimensional datasets for exploratory data analysis, low-dimensional representations of genetic similarity, which we refer to here as ancestry coordinates, are commonly computed and displayed as scatter plots. These visualizations are widely used to interpret population structure, and their geometry can shape how relationships between populations such as continuity, separation, and admixture are understood. Principal Component Analysis (PCA) [7] has traditionally been used for these visualizations [8–10]. But, as a linear method, it can struggle to capture localized patterns [11, 12] and loses detail and structure in the data when all but the first two or three components are discarded for visualization.

Nonlinear manifold learning methods such as t-SNE [13], UMAP [14], and PHATE [15] construct embeddings based on local neighborhoods defined at a chosen scale. Neighborhood embedding-based methods such as t-SNE and UMAP preserve these local groupings but do not preserve manifold distances within and between neighborhoods, and therefore often distort broader geometric relationships between populations [15, 16]. In particular, these methods tend to equalize apparent density across the embedding, so that clusters containing more samples occupy disproportionately more visual space—an effect viewers can misread as greater genetic separation or diversity [16, 17]. In population genetics, such distortions can obscure both nearby population relationships and larger-scale patterns such as geographic and admixture gradients, particularly when sampling gaps and population imbalance fragment continuous structure or exaggerate apparent separation between groups. Moreover, these representations risk oversimplifying the complexities of human diversity, falsely implying a lack of connection between groups. This highlights the need for visualization methods that more accurately reflect the structure of human genetic diversity without reinforcing misleading interpretations.

Methods based on diffusion geometry have been shown to better preserve such manifold structure by approximating diffusion processes on the underlying data manifold [15]. In particular, PHATE (Potential of Heat-diffusion for Affinity-based Trajectory Embedding) has been widely used to visualize high dimensional data, where it can capture continuous trajectories and multiscale relationships across a variety of biological datasets, ranging from single-cell transcriptomics and mass cytometry (CyTOF) profiling [15] to neuroscience [18]. However, the application of this method to population genetics remains underexplored.

Here, we present a PHATE-based approach to ancestry-coordinate construction that, when appropriately tuned, recovers and preserves continuous population structure. We show that commonly used ancestry coordinates can be systematically distorted by sampling gaps and population imbalance, and that naive use of PHATE can produce similar distortions. We focus on two construction choices that are especially consequential for population-genomic data: neighborhood scale and population-balanced fitting. Across large-scale genomic resources, including a combined HGDP and 1KGP reference panel (hereafter referred to as HGDP+1KGP) [19], UKBB, and AoU, we show that default ancestry coordinates can fragment continuous population structure or exaggerate dominant groups under realistic sampling conditions. Using a framework applicable at biobank scale, we further demonstrate that tuning neighborhood scale and fitting ancestry-coordinate methods on balanced subsets can restore interpretable trajectories that preserve geographic relationships and admixture gradients. To quantitatively evaluate embedding quality, we compare pairwise distances in the embedding space to distances derived from geographic sampling coordinates and inferred admixture proportions, which provide complementary references for continuous population structure. We additionally support these observations with synthetic experiments.

Together, these results provide a data-driven basis for constructing ancestry coordinates that better reflect continuous human genetic diversity and reduce the risk of over-interpreting disconnected clusters as discrete populations.

## 2 Results

### 2.1 A PHATE-Based Framework for Ancestry-Coordinate Construction

Diffusion-based methods provide a natural framework for representing the continuous and branching structure of human genetic variation. PHATE constructs a *k*-nearest neighbor graph linking each individual to their most similar neighbors, typically in principal-component space, then propagates these relationships across the graph by multi-step diffusion, summarizes the outcome as pairwise potential distances, and embeds these distances in two or three dimensions for visualization (Figure 1a; Supplementary Algorithm 1). Recovering accurate ancestry coordinates from this process is challenging, however: default parameter settings and preprocessing choices can introduce artifacts that distort or obscure the underlying population structure (Section 2.2). We therefore investigate two construction choices that together influence which features of population structure the resulting ancestry coordinates preserve.

**Fig. 1.**
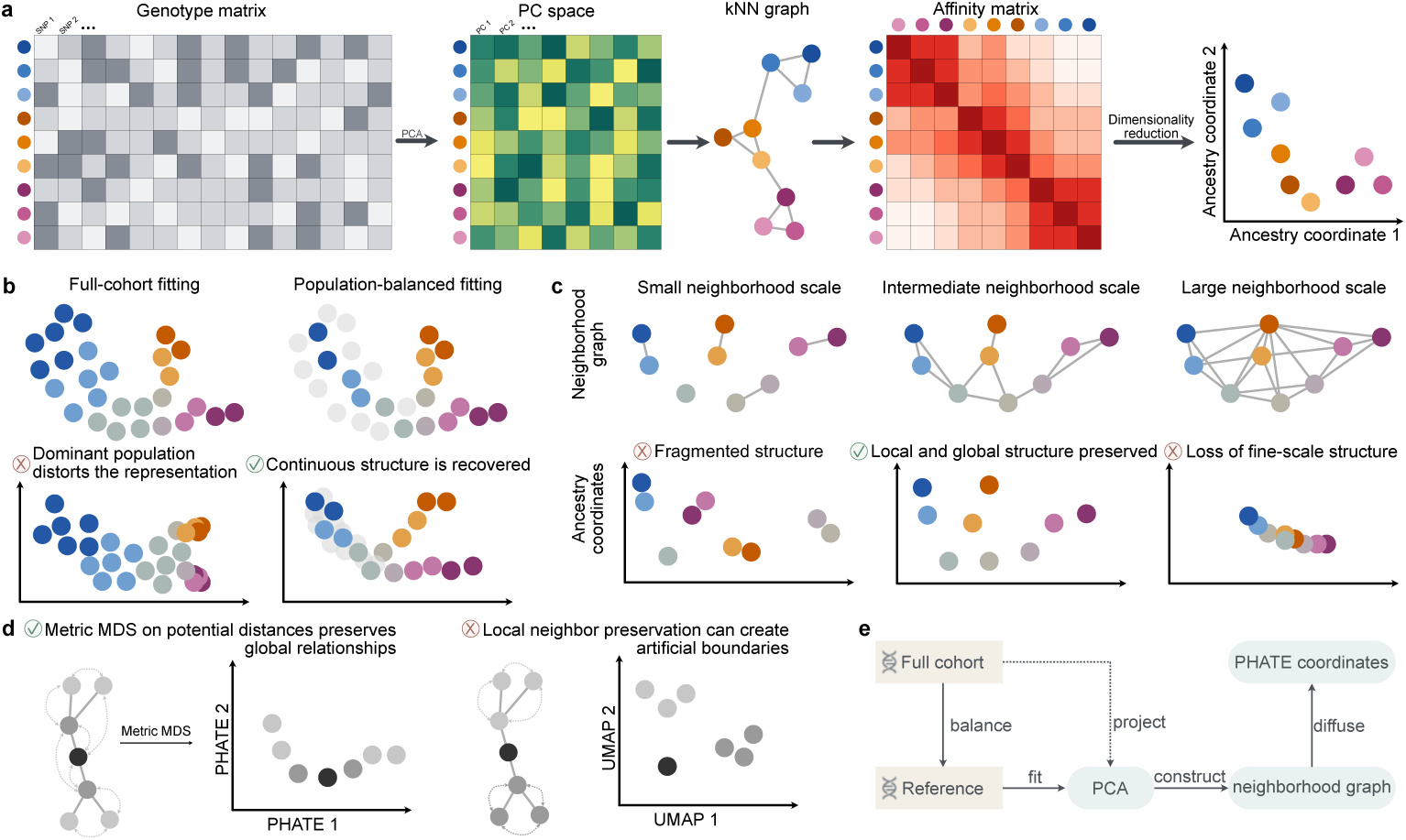
Conceptual framework for constructing PHATE-derived ancestry coordinates. (a) Schematic common to manifold learning methods explored in this paper (t-SNE, UMAP, and PHATE). Genotypes are represented as an individual-by-variant matrix, transformed into principal-component (PC) space, and used to construct a *k*-nearest-neighbor graph and affinity matrix. Manifold learning methods then use these affinities to generate two-dimensional ancestry coordinates. (b) Sampling composition influences ancestry-coordinate geometry: fitting on imbalanced cohorts can exaggerate dominant populations and distort continuous structure, whereas population-balanced fitting reduces sampling-density bias. (c) Neighborhood scale determines which aspects of structure are captured: small neighborhoods fragment continuous variation, intermediate neighborhoods preserve both local and global structure, and overly large neighborhoods compress fine-scale differences. (d) The embedding objective determines which relationships are preserved: PHATE fits all pairwise potential distances by metric multidimensional scaling (MDS), preserving global relationships between populations, whereas t-SNE and UMAP optimize the preservation of local neighborhoods only, which can introduce artificial boundaries between related groups. (e) Workflow at biobank scale: a population-balanced reference set is used to fit PCA and construct a neighborhood graph at an appropriate scale; PHATE is fitted on this reference graph. The remaining individuals can be projected into the resulting ancestry-coordinate space.

The first construction choice is sampling composition : fitting on a population-balanced subset rather than the full, potentially imbalanced cohort can reduce sampling-related distortions (Figure 1b). The second is the neighborhood scale, *k*: it determines the extent of genetic relationships captured by the neighborhood graph (Figure 1c). Although these choices also affect t-SNE and UMAP graph-construction steps (Algorithm 1 in Online Methods Section 4.1), PHATE’s embedding step, which fits all pairwise potential distances by metric multidimensional scaling rather than preserving only local neighborhoods, provides a means of capturing broader relationships between populations while retaining local genetic structure (Figure 1d). The theoretical and algorithmic results underlying these methods are presented in Supplementary Sections S1.1–S1.3. Building on this framework, we next evaluate the effects of sampling composition and neighborhood scale on the recovery of population structure across genomic datasets and synthetic benchmarks, following the practical workflow summarized in Figure 1e.

### 2.2 Default Ancestry Coordinates Reflect Sampling Gaps and Imbalances

Compared to data generated in controlled biological systems, human genetic datasets are subject to significant sociological constraints—such as participant consent, recruitment practices, logistical costs, and structural inequities—that systematically bias representation [20, 21]. These constraints result in geographic sampling gaps and population imbalances. We therefore first examine the globally diverse HGDP and 1KGP reference panels, which contain important gaps in worldwide representation, including limited representation from the Balkan Peninsula, Southeast Asia and Oceania. We then examined UKBB and AoU, two biobank cohorts from specific countries dominated by one or a few large groups, which reflect strong real-world population imbalance. For example, individuals who self-identify as British make up 95% of UKBB and around 50% of the AoU cohort self-identify as “White”. Using near-default parameter settings, we examined how commonly used ancestry coordinates represent population structure under these realistic, geographically incomplete, and imbalanced sampling regimes (Online Methods Sections 4.1 and 4.2).

Despite the broad geographic coverage of HGDP and 1KGP, all methods produced distorted representations of population structure (Figure 2). Ancestry continuity is absent from the t-SNE and UMAP representations, which instead fragment populations into detached clusters whose relative placement on the graph appears largely arbitrary, as illustrated by the Central and South Asian populations in the HGDP+1KGP UMAP embedding. The PHATE ancestry coordinates exhibit a distinct “floral” geometry, where sparsely sampled populations, including the under-represented Oceanian groups (Melanesian and Papuan), appear near the center of the visualization (the stigma), flanked by larger continental groups (petals).

**Fig. 2.**
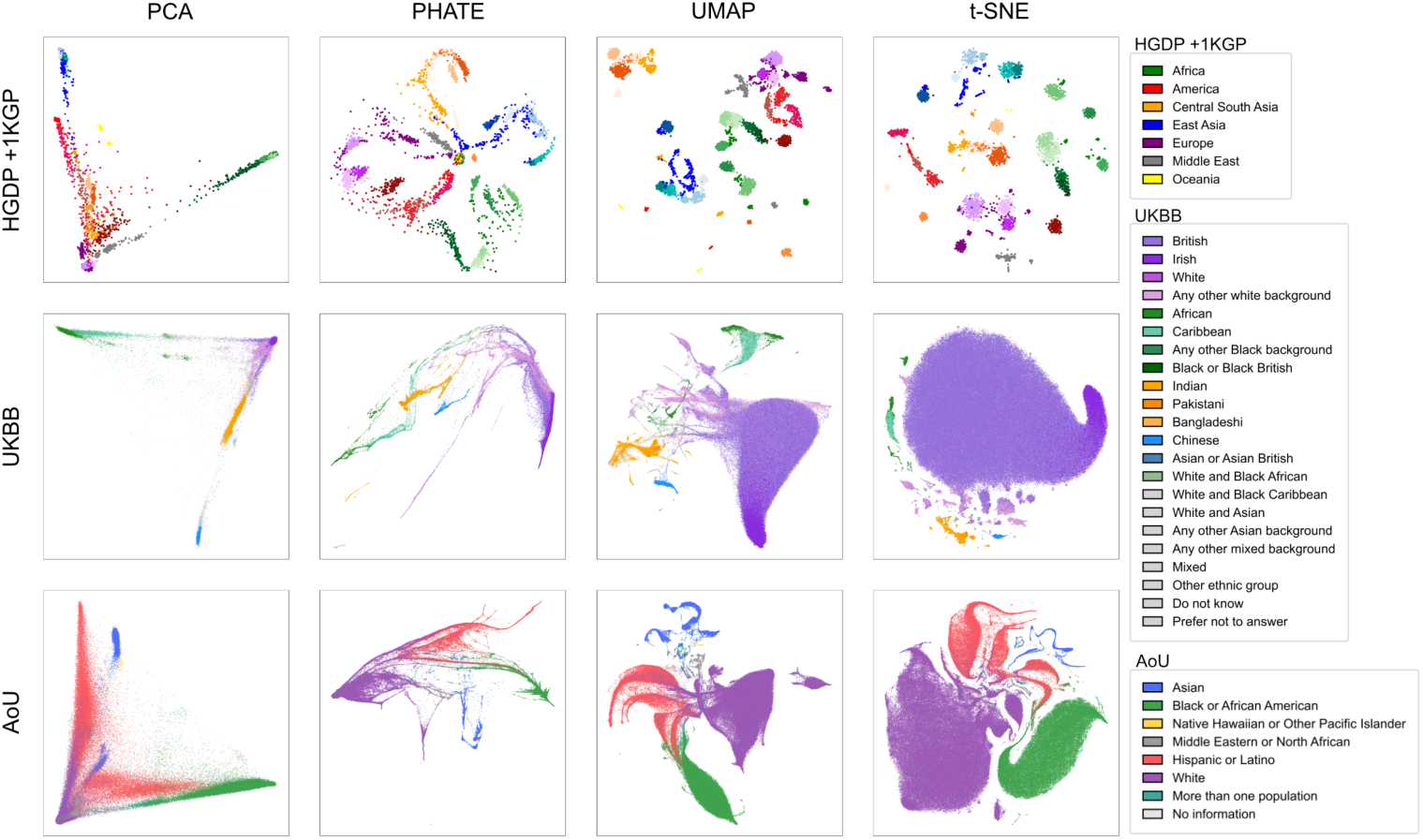
2D-PCA, t-SNE, UMAP and PHATE ancestry coordinates of the HGDP+1KGP (total *N* = 4,094), UKBB (total *N* = 486,748) and AoU (total *N* = 328,588) cohorts, using hyperparameters close to the defaults. For each dataset, we fitted the principal components on a balanced subset of unrelated individuals and projected the rest. The fitting sets contained *N* = 3,400 (HGDP+1KGP), *N* = 59,265 (UKBB), and *N* = 61,711 (AoU) individuals. See Section 4.1 for exact hyperparameter setting and Section 4.2 for data sources and preprocessing details.

We next examined UKBB and AoU, two large-scale biobanks characterized by substantial real-world population imbalance. For UKBB and AoU, the PHATE and UMAP representations appear as branching, hub-centered manifolds. Here, the pre-dominant population (British for UKBB and White for AoU) forms the central hub. Each branch corresponds to an admixture cline between the predominant population and another population represented in the cohort, including Indian, Caribbean and Chinese populations in UKBB. However, the visualization largely fails to capture admixture between non-dominant populations, with these individuals instead appearing collapsed onto overlapping regions of the embedding. Furthermore, for t-SNE and UMAP, a disproportionate fraction of the visualization is occupied by the predominant European population despite the lower genetic diversity of European populations relative to African and many admixed populations [4, 22]. Across all cohorts, the 2D-PCA embedding exhibits characteristic “horseshoe” shapes [12].

While our visualizations of the UKBB are consistent with previously reported UMAP plots [17], our AoU UMAP differs from the UMAP reported by the All of Us Research Program [6], for which the code and exact parameter settings were not released, limiting direct comparison. Together, these results show that default ancestry coordinates can fragment continuous structure, distort relationships between sparsely sampled populations, and over-emphasize dominant groups in large genomic datasets with uneven sampling.

### 2.3 Ancestry Structure Emerges from PHATE at Appropriate Neighborhood Scales

We next demonstrate the neighborhood-scale component of our ancestry coordinate framework, testing how this parameter shapes the representation of fine-scale versus broader population structure (Figure 1c). As introduced in Section 2.1, after PCA preprocessing, PHATE, UMAP, and t-SNE construct local neighborhoods among genetically similar individuals. Increasing the neighborhood scale, *k*, allows each individual to be connected to more genetic neighbors, shifting the resulting coordinates from very local similarity toward broader relationships spanning multiple populations. Here, we vary *k* in PHATE to test its effect on ancestry-coordinate structure.

We first theoretically characterized how neighborhood scale affects PHATE-based ancestry coordinates (Supplementary Section S1.5). When the neighborhood scale is too small, each individual is linked to only a handful of close neighbors, which can lead to a disconnected graph. We show that such graphs cause PHATE to treat individuals in different disconnected components as maximally distant, regardless of their true genetic differences, fragmenting populations into discrete clusters (Supplementary Section S1.5.1). When it is too large, nearly every individual becomes connected to nearly every other individual with similar affinity, so the graph can no longer distinguish nearby from distant individuals; the resulting affinities become nearly uniform and population structure collapses (Supplementary Section S1.5.2). These results imply that faithful ancestry coordinates can only emerge within an intermediate range of neighborhood scales.

We next verified whether this intermediate-scale regime could recover continuous ancestry structure in real genetic data without eliminating finer-scale population relationships, as expected from the theoretical characterization above. To test this empirically, we varied neighborhood scale *k* in PHATE applied to HGDP+1KGP (Extended Data Fig. 1). At small neighborhood scale (e.g., *k* = 5; Figure 2), the ancestry coordinates fragment into discrete clusters and exhibit the “floral” artifact, reflecting an overemphasis on local structure. As *k* increases, the ancestry coordinates undergo a transition toward continuous structure (Extended Data Fig. 1), progressively recovering relationships between populations. To characterize continuous ancestry variation, we estimated admixture proportions for HGDP+1KGP individuals and projected these components onto the coordinate space (Online Methods Section 4.2). Because HGDP+1KGP contains globally sampled populations with known geographic sampling locations, geographic distance provides an external reference for broad-scale population relationships (Online Methods Section 4.3).

At an intermediate neighborhood scale (*k* = 100; Figure 3), the ancestry coordinates preserve both broad east–west geographic structure and continuous admixture clines, including three-way European, African, and Indigenous American admixture (cf. Figure 2). Consistent with this visual pattern, geographic preservation increased as *k* was raised from near-default values to intermediate neighborhood scales (Supplementary Table S3). Notably, this increase in scale does not eliminate fine-scale structure, as individual populations remain clearly identifiable. We further show that the floral artifact cannot be resolved by adjusting other PHATE-specific settings alone, and that only sufficiently increasing *k* resolves it (Supplementary Section S1.7 and Supplementary Figure S1). Together, these results show that neighborhood scale determines whether ancestry structure appears fragmented or continuous, and that appropriate scaling enables simultaneous representation of both local and global population structure.

**Fig. 3.**
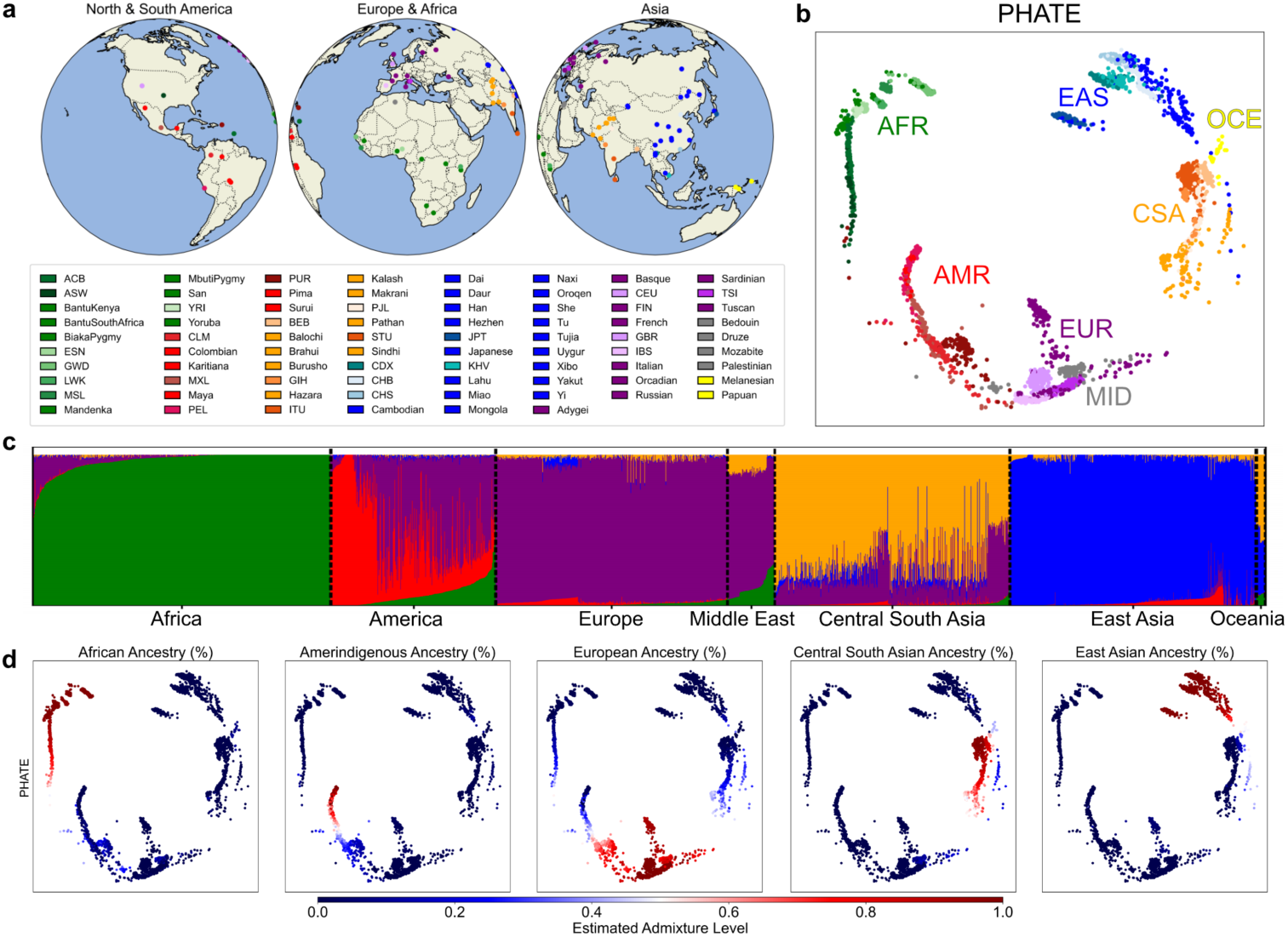
PHATE-derived ancestry coordinates recapitulate continuous population structure and admixture patterns in HGDP+1KGP. (a) Geographic distribution of sampled populations from HGDP+1KGP. Populations belonging to the same continent are similarly colored. Figure generated using the Cartopy Python package [23]. (b) PHATE ancestry coordinates (N=4,094) reveals a continuous organization aligned with known population geographic groupings. (c) Five-component ADMIXTURE ancestry proportions for HGDP+1KGP individuals in (b), grouped by population geographic region. ADMIXTURE components were interpreted according to the geographic populations in which they were most enriched (African, Amerindigenous, European, Central South Asian, and East Asian). (d) ADMIXTURE proportions projected onto PHATE space, illustrating the spatial organization of each ancestry component.

### 2.4 Synthetic Benchmarks Confirm Scale-Dependent Recovery of Latent Structure

Because real population-genetic datasets lack known ground truth, we next asked whether the scale-dependent changes observed in HGDP+1KGP reflect genuine recovery of latent continuous structure in the presence of sampling imbalance and population gaps.

To evaluate this, we generated a synthetic diffusion-limited aggregation (DLA) tree, a canonical hierarchical and tree-like geometry, with uneven sampling, and explicit gaps designed to mimic key challenges encountered in population-genetic datasets (Figure 4a, Online Methods Section 4.4, Supplementary Section S1.9.1). We quantified preservation by computing the correlation between embedded distances and ground truth distances (details in Section 4.3). PHATE recovered the global branching topology and continuous trajectories when an appropriate neighborhood scale is used (Figure 4b). When evaluated at their optimal parameter settings, most embedding methods achieve strong quantitative preservation of the ground-truth structure, with the exception of UMAP, which exhibits greater topological distortion (Figure 4c–d). However, the parameter regimes required to achieve optimal performance differ substantially across the different methods, consistent with the framework in Figure 1, where different methods implement neighborhood construction and embedding differently. Notably, t-SNE and UMAP require substantially larger neighborhoods to capture global branching, whereas diffusion-based methods (Diffusion Maps and PHATE) recover structure at more moderate scales. We found that PHATE produces visually coherent embeddings that preserve both global topology and local continuity across a range of *k* values explored (Figure 4c,e). Similar scale-dependent distortions can be observed across additional synthetic experiments, including Swiss roll, toroidal surface, and saddle surface examples (Supplementary Sections S1.9, S1.9.2, S1.9.3, and S1.9.4 and Supplementary Figure S9), which show that distortions arising under missing data and scale mismatch reflect a general property of embedding methods, rather than being specific to these datasets. Together, these findings show that the transition from fragmented to continuous structure observed in HGDP+1KGP reflects genuine recovery of latent geometry under uneven sampling and population gaps, rather than a dataset-specific artifact of the visualization.

**Fig. 4.**
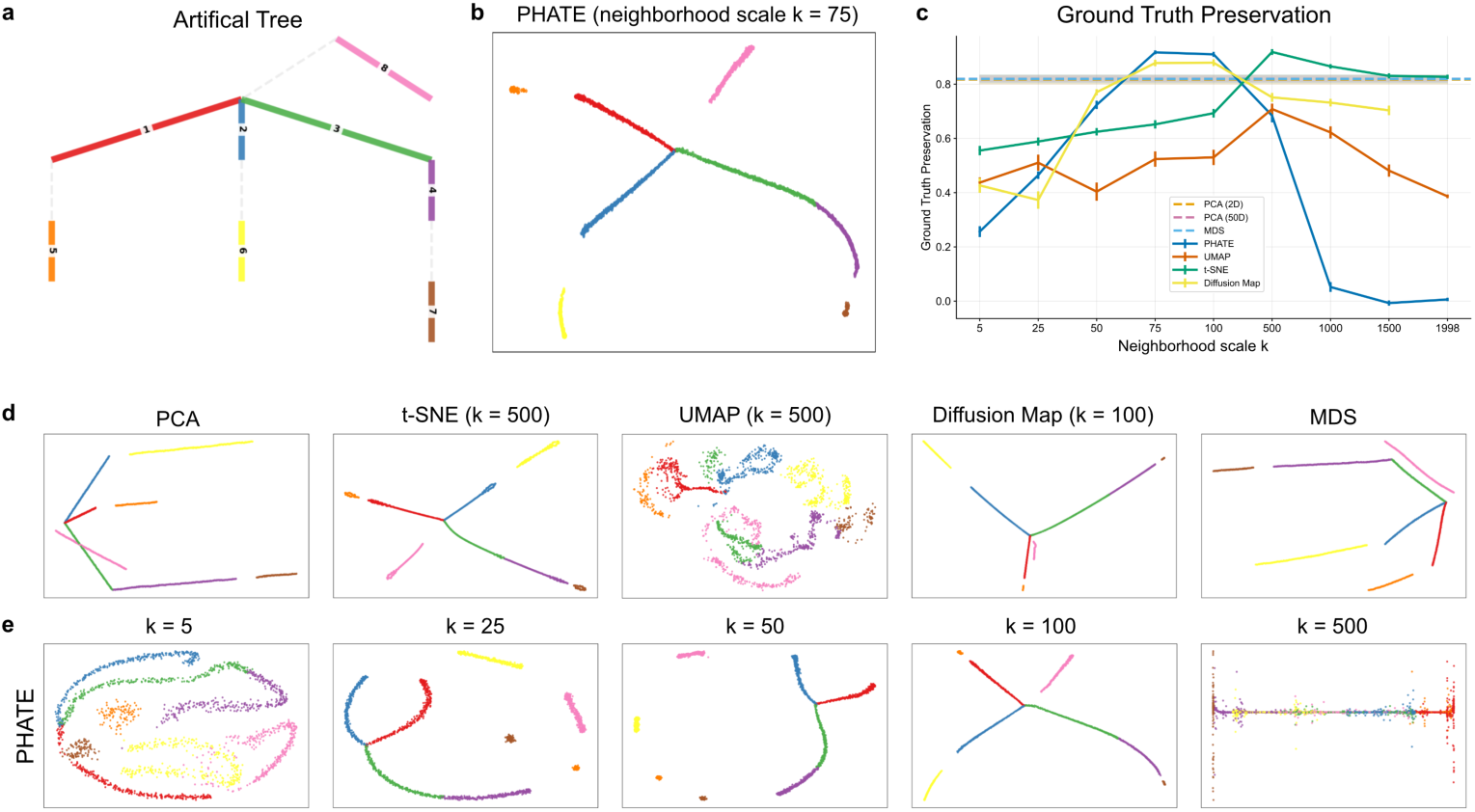
PHATE recovers continuous branching structure in synthetic DLA trees under appropriate neighborhood scales. a) Two-dimensional rendering of a synthetic DLA tree with color-coded branches, serving as ground-truth structure. b) PHATE embedding of the DLA tree (*k* = 75) accurately recovers the global branching topology and continuous trajectories of the ground truth. c) Quantitative assessment of ground-truth structure preservation, measured by Spearman correlation between embedded distances and corresponding geodesic distances along the complete latent DLA tree (Online Methods Section 4.3), across embedding methods and neighborhood sizes. d) Best-performing embeddings for PCA, t-SNE, UMAP, Diffusion Maps, and MDS, selected based on maximal ground-truth preservation. e) PHATE embeddings across a range of neighborhood scales (*k* = 5–500), highlighting degradation at overly small neighborhoods and stability at intermediate scales.

### 2.5 Population-Balanced PHATE Coordinates Preserve Admixture Structure in Large Biobank Datasets

Because the local neighborhoods used to construct ancestry coordinates are themselves shaped by sampling density (Figure 1b), we next asked whether analogous distortions could arise from population imbalance in real biobank cohorts, where sampling density varies substantially across ancestry groups. This question is motivated by a well-known observation in population genetics, where population imbalance is known to distort ancestry coordinates derived from PCA: over-sampled populations can dominate the visualization at the expense of less-sampled groups [11, 24]. To mitigate this effect, studies commonly fit PCA on a balanced subset of individuals from a cohort and then project the rest of the individuals. Nonlinear ancestry-coordinate methods such as t-SNE, UMAP and PHATE are then typically constructed from this projected PCA representation. However, these methods are often fitted on the full projected dataset [25, 26], thereby reintroducing imbalance during neighborhood graph construction. As shown in Figure 2, the population imbalance appears prominently in UMAP, t-SNE and PHATE ancestry coordinates of UKBB and AoU. For PHATE, we theoretically show that this distortion arises because the highly sampled populations dominate the neighborhood structure used to construct the coordinates, causing less-sampled populations and admixture trajectories to become compressed or disconnected (Supplementary Sections S1.1 and S1.6).

Because admixture provides a continuous measure of ancestry relationships in biobank cohorts, we asked whether population-balanced fitting combined with increased neighborhood scale could recover admixture structure in UKBB and AoU.

To reduce imbalance during neighborhood construction, we used the same population-balanced subsets used for PCA fitting (Figure 1e and Online Methods Section 4.2). Population-balanced fitting was constructed using pre-specified population labels, consistent with standard practice in population genetics. However, we found that a label-free sampling strategy produced similar ancestry coordinates in UKBB and AoU, suggesting that label-free balancing is also feasible (Supplementary Section S1.8 and Supplementary Figure S8).

To enable PHATE computation at biobank scale for the larger *k* values required for this analysis, we used an updated implementation (PHATE v2) based on randomized landmark selection, which improves scalability with respect to both sample size and *k* (Supplementary Section S1.10, Supplementary Figures S10 and S11).

Because the true geometry of population structure is not known in biobank data, we used inferred admixture proportions as an external reference for continuous ancestry variation. We quantified recovery of this structure using an admixture preservation metric, defined as the correlation between distances in the ancestry-coordinate space and distances derived from inferred admixture proportions (Online Methods Section 4.3). We computed this metric across increasing neighborhood scales and five random seeds. All methods exhibit poor admixture preservation at near-default parameter settings even after population-balanced fitting, but improve as neighborhood scale increases (Figure 5). Among the methods evaluated, PHATE achieved the strongest admixture preservation compared to t-SNE and UMAP for most neighborhood sizes. Balanced UKBB and AoU parameter sweeps further show that PHATE ancestry coordinates recover more continuous admixture structure at intermediate-to-large *k* and diffusion-time settings, whereas overly large scales again compress structure (Extended Data Figs. 2 and 3). These continuous PHATE trajectories correspond to gradients in inferred admixture composition (Supplementary Figures S6 and S7). By contrast, analyses on the full unbalanced UKBB cohort show persistent compression by the dominant sampled population across broad parameter sweeps for PHATE, t-SNE and UMAP (Supplementary Figures S2 and S3). Consistent comparisons across balanced UKBB and AoU also show that increasing neighborhood scale is insufficient for t-SNE and UMAP to recover the same admixture trajectories as PHATE, particularly among less-sampled ancestry groups (Supplementary Figures S4 and S5). Thus, the imbalance-driven distortion observed at near-default settings (Figure 2) is not resolved by increasing neighborhood scale alone in unbalanced data.

**Fig. 5.**
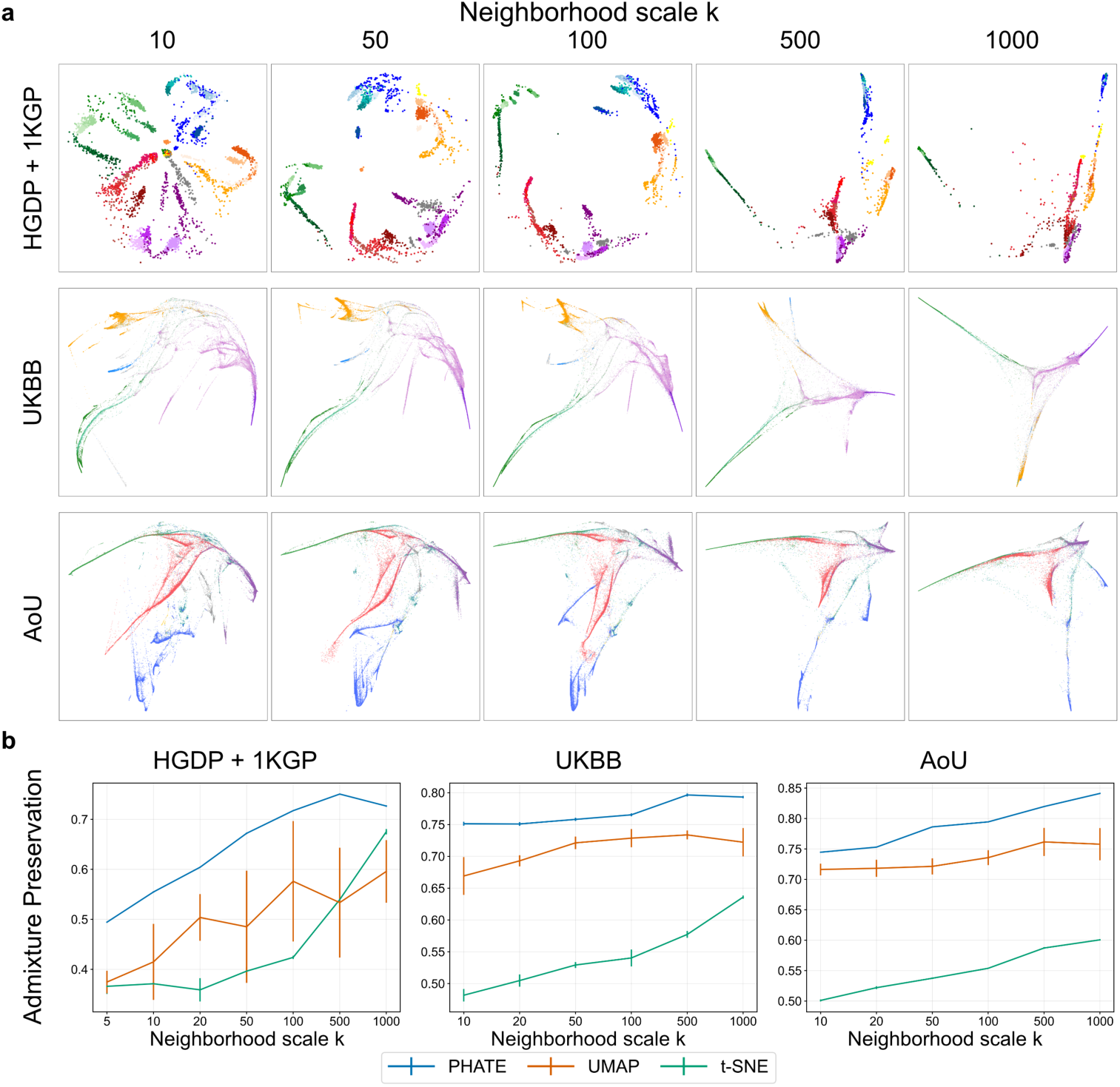
a) PHATE ancestry coordinates for HGDP+1KGP (*N* = 4,094), and for population-balanced fitting subsets of UKBB (*N* = 59,265) and AoU (*N* = 61,711), across increasing neighborhood scale *k* values. b) Admixture preservation for t-SNE, UMAP and PHATE across increasing neighborhood scales. Error bars represent standard deviation across random initializations. See Section 4.1 for exact hyperparameter setting. The full color legend is provided in Extended Data Fig. 4.

These results show that ancestry coordinates are shaped not only by underlying genetic structure, but also by the sampling distribution of the cohort itself, and that PHATE, balanced fitting, and increased *k* together enable recovery of interpretable continuous admixture trajectories in these large biobank datasets.

## 3 Discussion

Ancestry coordinates are not neutral summaries of genetic variation, and such visualizations can be misinterpreted even when produced with good intentions. Published nonlinear ancestry-coordinate plots often appear as detached blobs, stratified by population, including UMAP ancestry coordinates reported for gnomAD [27], AoU [6], UKBB [17, 25], and other genomic datasets [28]. These representations can give the false impression that there is no connection between the groups. In this study, we show that their geometry is shaped not only by underlying population structure, but also by methodological choices, including neighborhood scale, and by cohort composition, including sampling gaps and population imbalance. These choices affect whether relationships between populations are perceived as continuous, separated, or admixed, with direct consequences for the interpretation of ancestry gradients, migration history, and admixture.

In this context, appropriately tuned PHATE-derived ancestry coordinates recapitulate both geographic and admixture signals as trajectories that can be easily read from scatter plots. For the HGDP+1KGP experiments, broad geographic organization and admixture gradients are *simultaneously* visible in the PHATE representation, while individual populations remain distinguishable. This shows that continuity and fine-scale population differentiation are not mutually exclusive properties of diffusion-based ancestry coordinates. This behavior appears within an intermediate neighborhood-scale regime, where *k* controls the connectivity structure, providing a mechanistic interpretation of scale selection in population-genetic data. The scalable PHATE v2 implementation further shows that diffusion-based ancestry coordinates can be made feasible at biobank scale while preserving interpretable continuous structure.

Despite this, PHATE is also capable of producing problematic visualizations, as we demonstrate with default parameters (Figure 2). Comparable floral embeddings are also visible in PHATE plots previously published [26, 29]. To better preserve global population structure we significantly increased the neighborhood scale well beyond the default setting and computed our coordinates using the same balanced subset used for PCA computation. Both of these steps are essential, as increasing neighborhood size for unbalanced data resulted in ancestry coordinates with similarly distorted structure (see Supplementary Figure S2). These properties may be particularly important for admixed and underrepresented populations, where disconnected ancestry coordinates can obscure continuous ancestry gradients and compress relationships between groups.

A criticism of all visualization methods is that hyperparameters can be cherry-picked to produce ancestry coordinates that conform with preconceived biases [30]. Previous work has partially addressed this concern for t-SNE and UMAP in single-cell transcriptomics [31]; here, we address this in the context of population genetics data. We show that certain choices of hyperparameters and data sampling lead to failure modes, and that these choices can be evaluated by considering how they affect the topological properties of the diffusion operator and sampling imbalance (see Supplementary Sections S1.5 and S1.6). We evaluate ancestry coordinates using global structure preservation metrics, providing evidence of improved performance for both synthetic benchmarks and real biobank data. While we use hyperparameter sweeps to identify performant parameter regimes, establishing practical, theoretically motivated heuristics for hyperparameter selection is a natural and necessary direction for future work.

A limitation of applying these methods is that real biobank data lack known ground-truth population geometry. We therefore used two external references for continuous population structure: geographic distances, where sampling locations provide a meaningful proxy for broad-scale population relationships, and inferred admixture gradients, which capture shared ancestry and population connectivity. Although neither reference constitutes a definitive ground truth, together they provide biologically meaningful ways to assess whether ancestry coordinates align with broad patterns of human population history. Similarly, while population-balanced fitting based on pre-specified labels is standard practice in human genetics, such labels are imperfect and not always available. The label-free balancing results presented here suggest that this framework could generalize beyond label-defined population categories, motivating future work on automatic balancing and adaptive neighborhood selection. A further direction is to connect the diffusion process underlying PHATE to population-genetic theory. Applying PHATE to identity-by-descent (IBD) tract sharing or estimates of recent coalescence, and relating the resulting diffusion process to models of coalescing random walks on graphs over generations, could provide a theoretical basis for selecting neighborhood size *k* and diffusion time *t*.

Our results extend previous applications of PHATE by showing that neighborhood scale and population-balanced fitting are critical determinants of ancestry-coordinate geometry in population-genomic data. We showed that PHATE facilitates the visualization and interpretation of genetic diversity and population structure. Importantly, these analyses can be performed at biobank scale using our computationally efficient PHATE implementation. The resulting ancestry coordinates can distinguish between different populations while also showing the continuous variation between them, offering insights that are both scientifically meaningful and less likely to be misleading to the public. PHATE-based ancestry coordinates provide an alternative visualization framework, promoting a continuous view of ancestry [32]. These methods have the potential to help the field adopt representations of genetic data that better reflect human evolutionary history, in which global continuity and fine-scale population structure can be interpreted simultaneously.

## 4 Online Methods

### 4.1 Visualization Algorithms

We compared PCA [7], t-SNE [13, 33], UMAP [14], and PHATE [15] as low-dimensional ancestry-coordinate methods. 2D-PCA was used as a linear baseline, with PCs fitted on a subset of unrelated individuals and projected onto the full cohort following standard practice in population genetics. t-SNE and UMAP were used as nonlinear neighbor-preserving baselines, varying `perplexity` and `n_neighbors`, respectively. PHATE was used as a diffusion-based nonlinear embedding method. The main computational steps shared by these methods, and the differences in how PHATE, t-SNE and UMAP construct the final embeddings, are summarized in AlgorithmAlgorithm 1. For nonlinear methods, embeddings were computed from the top 20 projected PCs for each dataset. PCs were fitted on a subset for each dataset using PLINK v1.9 [34] and projected on the entire dataset. These were chosen by visual inspection, as they consistently showed population structure.

#### Algorithm 1

Manifold-learning-based ancestry-coordinate construction pipeline. PHATE and t-SNE/UMAP diverge mainly in the embedding step (4a vs. 4b). Exact kernels, normalizations, and objectives are given in Supplementary Sections S1.1–S1.3.

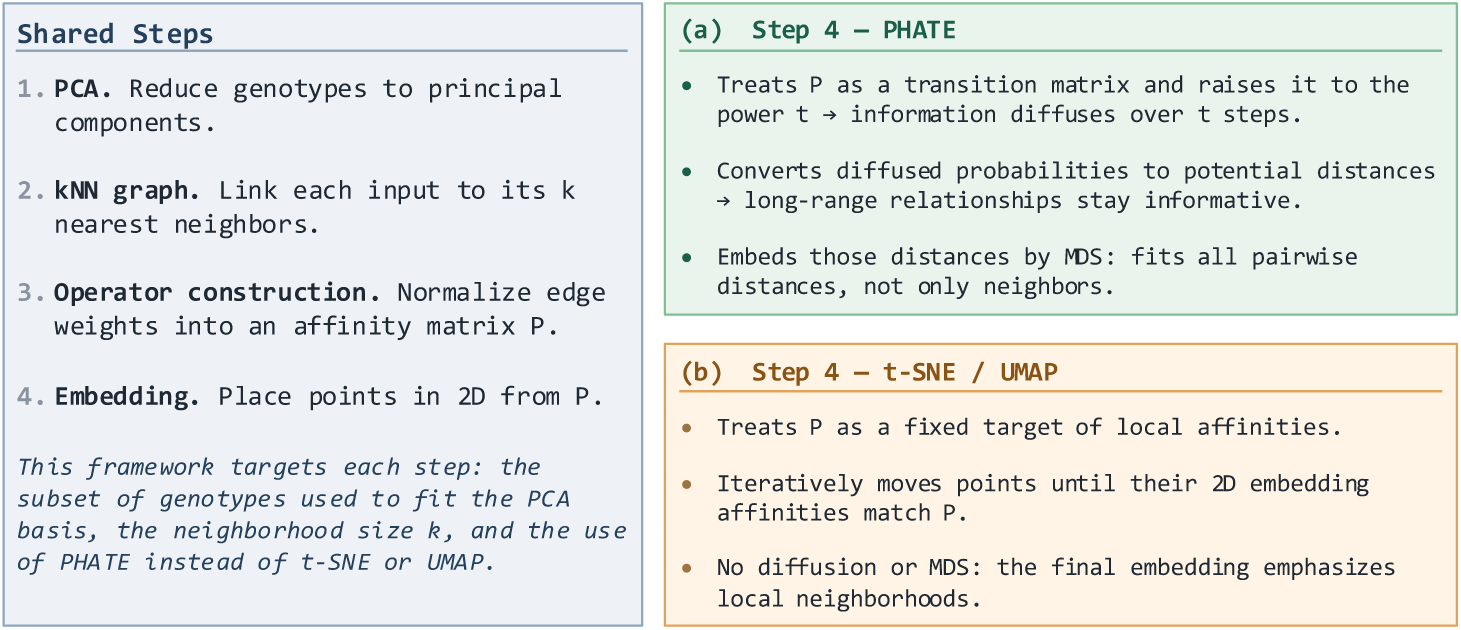

All three nonlinear methods share an affinity-construction step. Let *X* = *{x*_1_,*…, x_N_}* denote the genetic PC representations of *N* individuals. Pairwise distances among these representations define a matrix of affinities **K** from a similarity kernel, built from a per-individual bandwidth *ω_i_*(*k*), schematically

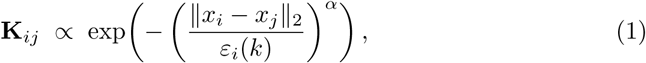

which is normalized into a method-specific operator **P** that encodes the relationships preserved in the low-dimensional ancestry coordinates (exact kernels and normalizations in Supplementary Sections S1.1–S1.3). The neighborhood term sets this bandwidth: in PHATE the `KNN` parameter fixes *ω_i_*(*k*) as the distance from *x_i_* to its *k*-th nearest neighbor, and in t-SNE and UMAP the analogous bandwidth is set by `perplexity` and `n_neighbors`, respectively. We refer to this shared neighborhood-size parameter as *k* throughout the main text, following the notation used in the original PHATE and UMAP papers themselves; each method’s software instead exposes it under its own argument name (`knn` in the PHATE package [15], `n neighbors` in `umap-learn` [14], and `perplexity` in `openTSNE` [33]). Widening *ω_i_*(*k*) raises the off-diagonal affinities and so increases the number of pairwise connections represented by **K** and **P**. While `KNN` and `n_neighbors` are roughly comparable, neither corresponds exactly to `perplexity`, though `n_neighbors` is often approximated as 3*↔*`perplexity` [35]. In PHATE, increasing diffusion time *t* and decreasing kernel decay have an effect similar to increasing `KNN`, but these parameters are not interchangeable (see Supplementary Section S1.7).

The main hyperparameter settings used for the default and neighborhood-scale experiments are summarized below. For Figure 2, all ancestry coordinates were generated using default or commonly used parameter settings. PHATE was run with `KNN` = 5, decay = 40, and *ε* = 1; t-SNE used a perplexity of 33; and UMAP used `n_neighbors` = 10. For HGDP+1KGP, PHATE was run with *t* = 10 and UMAP with `min_dist` = 1.0, whereas for UK Biobank and All of Us, PHATE used *t* = 100 and UMAP used `min_dist` = 0.5. For Figure 5 and Supplementary Table S3, PHATE was run with decay = 40 and *ε* = 1, and UMAP used `min_dist` = 0.5. For HGDP+1KGP, PHATE was run with *t* = 25; for UK Biobank and All of Us, PHATE used *t* = 100. Further PHATE details are provided in Supplementary Sections S1.1 and S1.10.

### 4.2 Data Sources and Preprocessing

We analyzed three large-scale human genomic resources: the combined Human Genome Diversity Project and 1000 Genomes Project reference panel (HGDP+1KGP), the UK Biobank (UKBB), and the All of Us (AoU) Research Program. Genotypes were restricted to high-quality autosomal single-nucleotide polymorphisms (SNPs), with standard filtering for missingness, minor allele frequency, and linkage disequilibrium (see Supplementary Section S1.4). Genotypes were harmonized across datasets by aligning alleles to the HGDP+1KGP reference panel, resolving strand inconsistencies, and restricting analyses to a common set of high-quality SNPs shared across cohorts.

For HGDP+1KGP, a maximal subset of 3,400 unrelated individuals was selected using KING relatedness [36]. Admixture proportions were estimated on the HGDP+1KGP reference panel using ADMIXTURE [37] and Neural ADMIXTURE version 1.6.7 [38], for computational efficiency on biobank cohorts (Online Methods Section 4.3). For within-cohort visualization in UKBB and AoU, ancestry coordinates were computed on subsets from each biobank cohort to control for population imbalance. In UKBB, a subset of 59,265 unrelated individuals was used, subsampling to 10,000 White British and 5,000 Irish individuals, following [5] (see Supplementary Section S1.4.2). For AoU, a subset of 61,711 unrelated individuals was used, where we subsampled to 10,000 for each predominant group: “White”, “Hispanic or Latino” and “Black or African American” (Supplementary Section S1.4.3). For cross-cohort analyses, we first computed PCA and estimated admixture models using the HGDP+1KGP reference panel (Supplementary Section S1.4.1), and all UKBB and AoU samples were then projected onto these reference-derived PCA axes and admixture components. Detailed preprocessing, quality control, and harmonization procedures for each dataset are provided in Supplementary Section S1.4.

### 4.3 Evaluation Metrics

For genomics datasets, we define metrics tailored to population genetics, based on geographic and admixture information. We used the following metrics:

1. **Geographic preservation**: Spearman correlation between embedded distances and the Haversine (Great-circle) distance using the longitude and latitude of the samples. This metric has been used before [19, 39]. We excluded samples from the Americas, because recent admixture and colonial-era demographic history decouple present-day sampling location from the geographic origins of ancestry components (see Supplementary Table S1).
2. **Admixture preservation**: Spearman correlation between embedded distances and geodesic distance using admixture ratios as coordinates. We report the average correlation varying between 2–9 components. For HGDP+1KGP, we estimated admixture ratios using ADMIXTURE [37]. For UKBB and AoU, we used neural admixture version 1.6.7 to estimate admixture proportions by fitting the model on HGDP+1KGP and projecting each biobank cohort onto the resulting reference components [38].
3. **Ground-truth preservation** (synthetic data): Spearman correlation between embedded distances and geodesic distances along the complete latent structure before gap exclusion (Online Methods Section 4.4).

For UKBB and AoU, we subsampled 5,000 individuals and computed the correlation metrics on these, instead of the full dataset. Metrics were computed separately for each biobank, embedding method, neighborhood-scale parameter, and random seed. We compared PCA, t-SNE, UMAP, and PHATE across the analyzed cohorts, varying perplexity for t-SNE, `n_neighbors` for UMAP, and `KNN` for PHATE; other hyperparameters, including PHATE diffusion time *t* and UMAP `min_dist`, were fixed as described in Section 4.1.

### 4.4 Synthetic Data

We generated synthetic datasets to evaluate whether embedding methods recover continuous population structure when the ground truth is known. Our primary synthetic benchmark was a graph-conditioned diffusion-limited aggregation (DLA) tree, a hierarchical and tree-like geometry in which samples were generated along the branches of a predefined directed tree rather than from an unconstrained recursive attachment process [15]. This design allowed us to control features that are difficult to isolate in real genetic data, including branching topology, uneven sampling across branches, and missing populations.

To model sampling gaps, selected branches or branch segments were generated as part of the latent tree but excluded from the observed dataset. Gaussian noise was then added to retained samples to mimic measurement variability. Ground-truth relationships were defined by geodesic distances along the complete latent tree before gap exclusion, providing a controlled reference for evaluating whether each embedding method preserved intrinsic structure rather than only ambient Euclidean proximity. Full details of the DLA construction, gap generation, noise model, and ground-truth distance calculation are provided in Supplementary Section S1.9.1.

### 4.5 Scalable implementation of PHATE

To enable PHATE analyses at biobank scale, we used a scalable landmark-based implementation. Instead of relying on the spectral approach based on randomized Singular Value Decomposition (SVD) and iterative MiniBatchKMeans clustering, our implementation directly selects *n*_landmark_ points uniformly at random as landmarks. Each sample is then assigned to its nearest landmark by computing pairwise Euclidean distances. Landmarks are sampled uniformly without replacement; we used *n*_landmark_ = 10,000 for both UKBB and AoU. This eliminates the costly eigendecomposition and iterative clustering steps, reducing the overall complexity from *O*(*n*^2^*k*+*nkt*) (dominated by SVD and KMeans iterations) to a simpler *O*(*n·n*_landmark_). As a result, the runtime grows almost linearly with both *n* and *n*_landmark_, making the approach significantly faster and more scalable for large datasets, while preserving the qualitative structure of the embeddings. Detailed implementation and benchmarking results are provided in Supplementary Section S1.10.

## Data availability

The HGDP+1KGP reference panel used in this study was obtained from the gnomAD v3.1.2 release and is available through the gnomAD download portal (https://gnomad.broadinstitute.org/downloads). UK Biobank data are available to approved researchers through the UK Biobank Access Management System (https://www.ukbiobank.ac.uk/use-our-data/apply-for-access/); these analyses were conducted under UK Biobank application 20168. The All of Us data analyzed here are from the All of Us Research Program Controlled Tier Dataset v7. Participant-level All of Us data cannot be redistributed by registered users under the All of Us Data and Statistics Dissemination Policy (https://support.researchallofus.org/hc/en-us/articles/22346276580372-Data-and-Statistics-Dissemination-Policy). Aggregate information is available from the All of Us Research Program website (https://www.researchallofus.org/), and eligible researchers may register for access through the Researcher Workbench (https://www.researchallofus.org/register/).

## Code availability

Code for the preprocessing pipeline, including PCA computation and admixture estimation is available at github.com/MattScicluna/manifold_genetics. PHATE code (including random landmarking) is available at github.com/KrishnaswamyLab/PHATE. Hyperparameter sweeps were performed using the `manylatents` software package github.com/latent-reasoning-works/manylatents. All of Us-specific analysis code is available in the All of Us Researcher Workbench 2.0 workspace “PHATE-derived ancestry coordinates capture continuous population structure” (https://workbench.verily.com/workspaces/phate-derived-ancestry-coordinates-capture-continuous-populatio). Workspace access is available upon request to researchers with the required All of Us data access permissions.

## Ethics approval and consent to participate

This study was approved by the Research Ethics Board of the Montreal Heart Institute (MHI; project 2026-3546). All genomic data were accessed through established biobank access procedures, following the required data access approvals and completion of relevant research ethics training. Informed consent was obtained from participants by the respective biobanks in accordance with their approved protocols.

## Author contributions

Author contributions are described following the CRediT taxonomy. **Conceptualization:** M.S., G.W., S.K., J.G.H. **Methodology:** M.S., S.N., C.M.V.C., J.C.G., G.W., S.K., S.L., J.G.H., K.R.M. **Software:** M.S., S.N., C.M.V.C., J.F.R., M.E.K. **Formal analysis:** M.S., S.N. **Investigation / experiments:** M.S., S.N., J.C.G., M.E.K. **Data curation:** M.S., J.C.G., R.P. **Visualization / figures:** M.S., S.N. **Writing – original draft:** M.S., S.N. **Writing – review & editing:** M.S., S.N., C.M.V.C., J.C.G., R.P., G.W., S.K., J.G.H., K.R.M., S.L. **Supervision:** S.L., G.W., S.K., J.G.H.

## Declaration of Use of Artificial Intelligence (AI)-assisted technologies

During the preparation of this work, the authors used large language models to improve the readability and grammar of this manuscript, as well as assisting with code generation. After using these tools, the authors reviewed and edited the manuscript and associated code and take full responsibility for the content of the published article. No AI tools were used for data interpretation or for the generation of scientific conclusions.

## Acknowledgements

We thank John Novembre, Alex Diaz-Papkovich, Fatima Mostefai, Dhananjay Bhaskar, Charles Xu, Chen Liu, and members of the MHI-OMICS group, notably Ndikumasabo Ratzy-Chris, for valuable discussions regarding this work.

This work was made possible through computational resources provided by the Digital Research Alliance of Canada and by Mila (mila.quebec). We also thank the participants of the 1KGP, HGDP, UKBB and the AoU biobank for their contributions which have made this research possible.

## 5 Extended Data Figures

**Extended Data Fig. 1.**
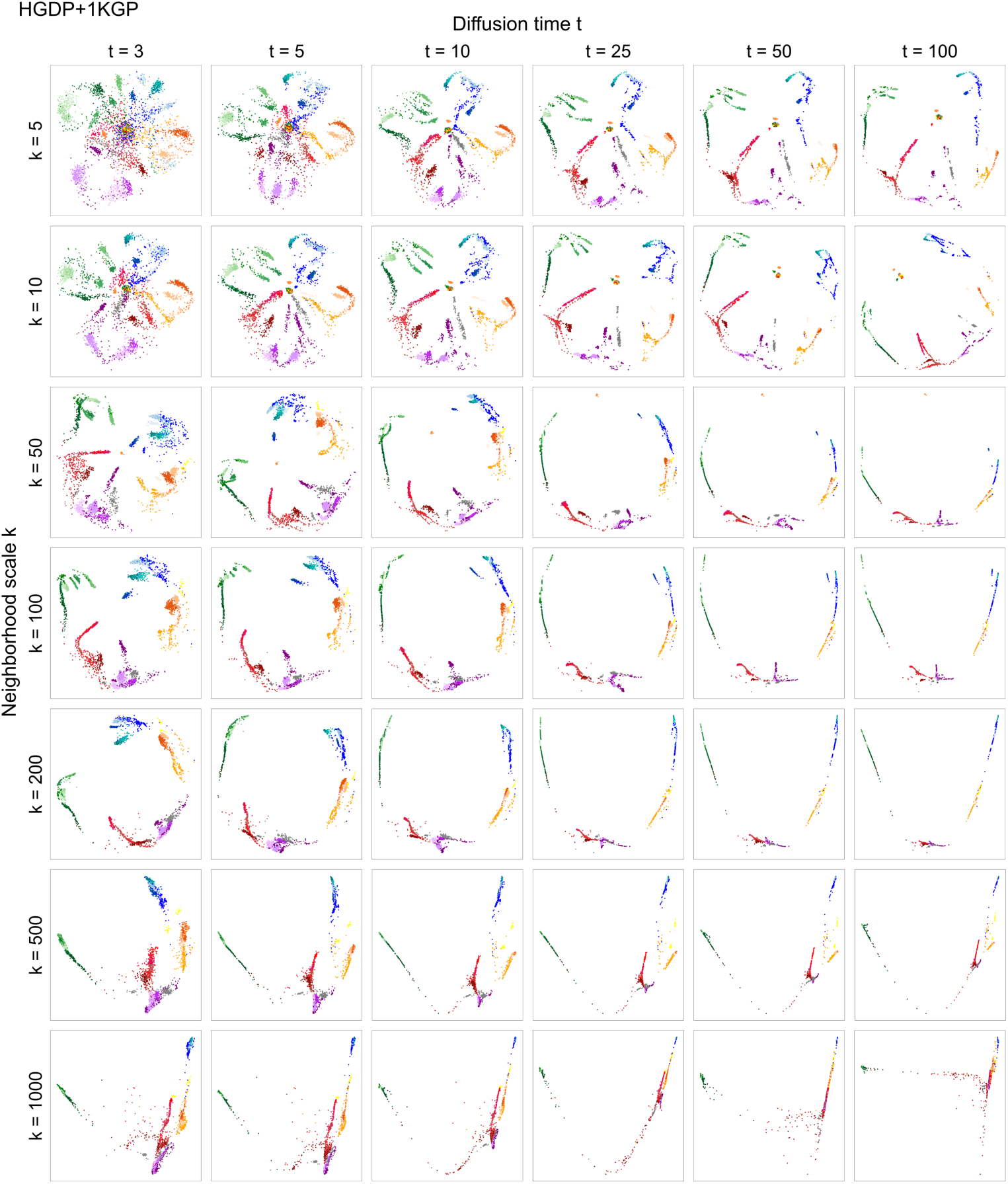
PHATE fitted to HGDP+1KGP (*N* = 4,094) varying neighborhood scale *k* and diffusion time *t*. All other parameters are set to default.

**Extended Data Fig. 2.**
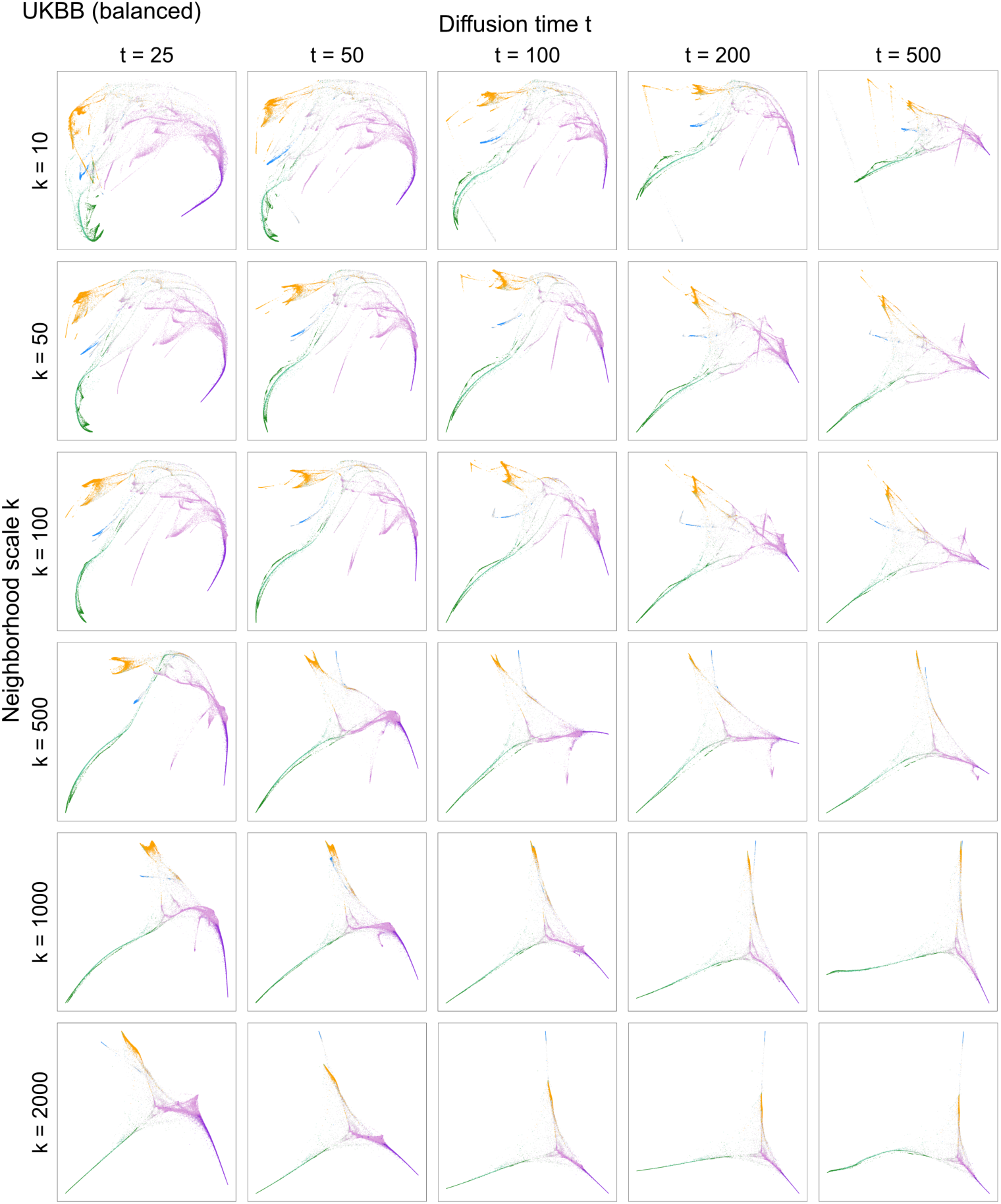
PHATE fitted to balanced UKBB (*N* = 59,265) varying neighborhood scale *k* and diffusion time *t*. All other parameters are set to default.

**Extended Data Fig. 3.**
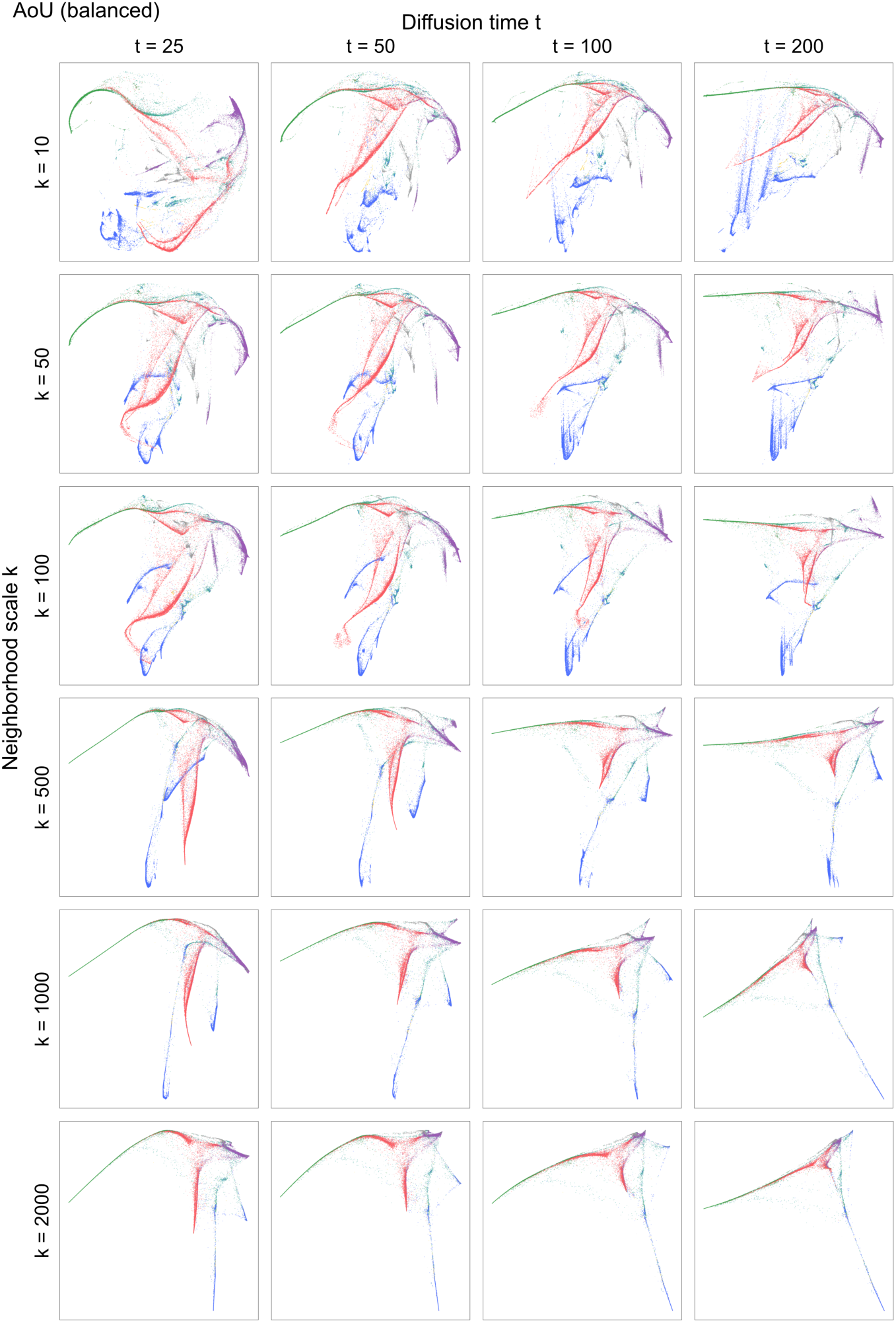
PHATE fitted to balanced AoU (*N* = 61,711) varying neighborhood scale *k* and diffusion time *t*. All other parameters are set to default.

**Extended Data Fig. 4.**
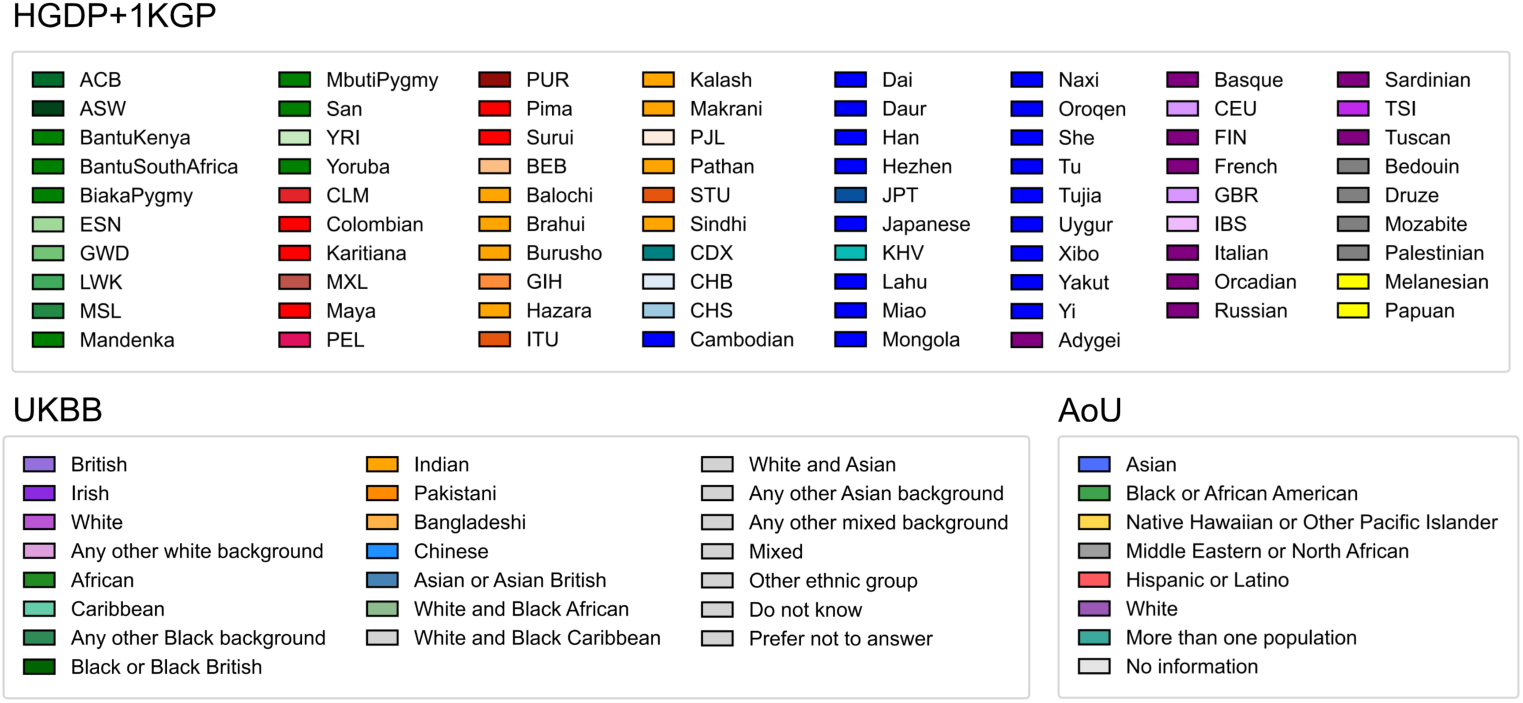
Color legend for ancestry and population labels across HGDP+1KGP, UK Biobank, and All of Us Biobank. (Extended Data Figs. 1–3)

