## Supplementary text, figures and tables for "PHATE-Derived Ancestry Coordinates Capture Continuous Population Structure in Human Genomic Data"

### S1 Supplementary Text

#### S1.1 PHATE algorithm and parameterization

In this section, we summarize the components of the PHATE algorithm that are relevant to our analysis of neighborhood scale and sampling effects in ancestry embeddings. For a detailed treatment, see the original paper [15].

PHATE is a manifold learning algorithm that combines ideas from diffusion geometry and information theory to denoise and visualize both global and local structure in high-dimensional data. The key assumption underlying PHATE is that data lie on a manifold with gradual transitions between samples that form coherent paths in a limited number of directions. These paths can branch, creating tree-like or trajectory-like structures.

The first step of PHATE is to construct a Markov transition matrix that encodes local sample affinities. Let $x_1, \ldots, x_N = \mathcal{X}$ be the dataset. The diffusion operator $\mathbf{P}$ is constructed by row-normalizing a matrix of kernel values $k$:

$$[\mathbf{P}]_{(x,y)} = \frac{k(x,y)}{\sum_{z \in \mathcal{X}} k(x,z)}, \quad x, y \in \mathcal{X}. \tag{E1}$$

PHATE uses an $\alpha$-decaying kernel:

$$k_{k,\alpha}(x,y) = \frac{1}{2} \exp\left(-\left(\frac{\| x-y \|_2}{\varepsilon_k(x)}\right)^{\alpha}\right) + \frac{1}{2} \exp\left(-\left(\frac{\| x-y \|_2}{\varepsilon_k(y)}\right)^{\alpha}\right), \tag{E2}$$

where $\varepsilon_k(x)$ is the distance from $x$ to its $k$th nearest neighbor. The two key hyperparameters here are $k$ (also referred to as `KNN` in the PHATE package [15]), which determines the size of local neighborhoods, and $\alpha$, which controls how sharply the kernel decays with distance.

Informally, increasing $\alpha$ leads to more constant kernel values around $(-\varepsilon_k(x), \varepsilon_k(x))$, and lighter tails. Increasing `KNN` increases $\varepsilon_k(x)$. In Section S1.5, we show how setting $k$ inappropriately can lead to distorted embeddings.

PHATE denoises the data by applying the diffusion operator $\mathbf{P}$ repeatedly: the $t$-step diffusion operator $\mathbf{P}^t$ corresponds to a random walk of length $t$, and the parameter $t$ governs the extent of the denoising. For a point $x$, we denote the $x$-th row of $\mathbf{P}^t$ as $p_x^t = \delta_x \mathbf{P}^t$ (where $\delta_x$ is the Dirac delta centered at $x$). For any $z \in \mathcal{X}$, we denote $p_x^t(z)$ as the entry of $p_x^t$ corresponding to $z$, suppressing $t$ where convenient.

The final distance used for embedding is the potential distance:

$$\mathcal{D}(x,y) = \| U_x - U_y \|_2, \tag{E3}$$

where $U_x = -\log(p_x^t)$. That is, PHATE embeds the log-transformed diffusion probabilities rather than the raw distances or transition probabilities. This potential distance reflects how easily two samples can reach similar neighborhoods via multi-step random walks. It emphasizes shared structure while down-weighting noise and outliers, offering a more faithful representation of manifold geometry than Euclidean distance or standard diffusion distances. We note that the PHATE implementation also allows for alternative informational distances, which we do not discuss here.

In our analysis, two aspects of this are particularly important: the neighborhood size parameter $k$, and the diffusion operator $\mathbf{P}^t$ which propagates information.

Two implementation details will be significant for our later proofs (Section S1.5):

1. PHATE sets kernel values below a certain threshold to 0. We define $p_{\min}$ as the minimum non-zero value of the diffusion operator $\mathbf{P}$.
2. A small constant $\epsilon$ is added to all entries of $\mathbf{P}$ to prevent numerical instability.

PHATE computes the final low-dimensional embedding by applying classical multidimensional scaling (MDS) followed by metric MDS to the matrix of potential distances.

#### S1.2 t-SNE algorithm and parameterization

In this section we summarize the components of t-SNE [13], as implemented in the `openTSNE` package [33], relevant to our neighborhood-scale analysis; for a complete treatment see the original paper. Like PHATE, t-SNE row-normalizes a local kernel, but then symmetrizes these per-point affinities into a single joint distribution and embeds the data using only these one-step affinities.

Let $\mathcal{X} = \{x_1, \ldots, x_N\}$ be the dataset. t-SNE uses a Gaussian kernel $\mathbf{K}_{ij} = \exp(-\parallel x_i - x_j \parallel_2^2 /2\sigma_i^2)$, which it row-normalizes and symmetrizes into a joint distribution $\mathbf{P}$:

$$\mathbf{P}_{j|i} = \frac{\mathbf{K}_{ij}}{\sum_{l \neq i} \mathbf{K}_{il}}, \qquad \mathbf{P}_{ij} = \frac{\mathbf{P}_{i|j} + \mathbf{P}_{j|i}}{2N}, \tag{E4}$$

with $\mathbf{P}_{ii} = 0$ and $\sum_{i,j} \mathbf{P}_{ij} = 1$. As in PHATE (Equation E1), $\mathbf{P}$ is derived from a kernel by normalization, but the row-normalized per-point conditionals $\mathbf{P}_{\cdot|i}$ are here symmetrized into a single joint distribution (summing to one over all pairs) rather than kept row-stochastic as in PHATE. The per-point bandwidth $\sigma_i$ is the neighborhood-scale parameter: it is not set directly but calibrated for each point so that the perplexity of $\mathbf{P}_{\cdot|i}$ matches a user-specified target. Increasing perplexity widens $\sigma_i$ and the effective neighborhood—the t-SNE analogue of `KNN` in PHATE [16, 35]—shifting the embedding toward larger-scale structure.

The embedding $\{y_1, \ldots, y_N\}$ is fitted by minimizing the Kullback–Leibler divergence between $\mathbf{P}$ and a low-dimensional affinity $\mathbf{Q}$, built from a heavy-tailed Student-$t$ kernel $\mathbf{Q}_{ij} \propto (1+ \parallel y_i - y_j \parallel_2^2)^{-1}$ and normalized so that $\sum_{i,j} \mathbf{Q}_{ij} = 1$:

$$\min_{y_1,\ldots,y_N} \sum_{i,j} \mathbf{P}_{ij} \log \frac{\mathbf{P}_{ij}}{\mathbf{Q}_{ij}}. \tag{E5}$$

Because the divergence heavily penalizes placing a high-affinity pair far apart in the embedding, t-SNE prioritizes local neighborhood structure over large-scale geometry. The objective is non-convex and is minimized by gradient descent.

#### S1.3 UMAP algorithm and parameterization

UMAP [14] constructs a similar neighbor graph but frames embedding as preserving a fuzzy topological representation of the data.

Let $\mathcal{X} = \{x_1, \ldots, x_N\}$ and fix a neighborhood size $k$ (the `n_neighbors` parameter, default 15; the UMAP analogue of `KNN`). For each $x_i$, let $\mathcal{N}_i$ be its $k$ nearest neighbors and $\rho_i$ the distance to its nearest neighbor. A per-point bandwidth $\sigma_i$ is calibrated from $k$ via $\sum_{x_j \in \mathcal{N}_i} \exp(-\max(0, \| x_i - x_j \|_2 - \rho_i)/\sigma_i) = \log_2 k$, giving the directed affinities

$$\mathbf{P}_{j|i} = \mathbf{K}_{ij} = \exp\left(-\frac{\max(0, \| x_i - x_j \|_2 - \rho_i)}{\sigma_i}\right), \tag{E6}$$

symmetrized by a fuzzy union $\mathbf{P}_{ij} = \mathbf{P}_{j|i} + \mathbf{P}_{i|j} - \mathbf{P}_{j|i}\mathbf{P}_{i|j}$ (and, unlike t-SNE, not normalized to sum to one). Increasing `n_neighbors` widens $\sigma_i$ and thus $\mathcal{N}_i$. Because $\mathbf{P}_{j|i} = 0$ whenever $x_j \notin \mathcal{N}_i$, these affinities are supported entirely on the (symmetrized) $k$-nearest-neighbor graph: every non-neighbor pair receives zero membership and is therefore undifferentiated, in contrast to PHATE's multi-step diffusion operator $\mathbf{P}^t$, which relates non-adjacent individuals through intermediate paths.

The embedding $\{y_1, \ldots, y_N\}$ is fitted by minimizing the fuzzy-set cross-entropy between $\mathbf{P}$ and a low-dimensional membership $\mathbf{Q}_{ij} = (1+a \| y_i - y_j \|_2^{2b})^{-1}$, where the parameters $a$ and $b$ are determined from the `min_dist` hyperparameter (default 0.1):

$$\min_{y_1,\ldots,y_N} \; -\sum_{i \neq j} \Big[\mathbf{P}_{ij} \log \mathbf{Q}_{ij} + (1 - \mathbf{P}_{ij}) \log(1 - \mathbf{Q}_{ij})\Big]. \tag{E7}$$

Like t-SNE, this objective emphasizes local neighborhoods over global geometry. It is minimized by stochastic gradient descent, with the repulsive (second) term approximated by negative sampling; this approximation causes the effective objective to differ somewhat from the stated cross-entropy [40].

#### S1.4 Data preprocessing and cohort harmonization

##### S1.4.1 1KGP and HGDP

We obtained the combined HGDP+1KGP data from the gnomAD v3.1.2 release [27]. Variants were restricted to autosomal single-nucleotide polymorphisms (SNPs) with `PASS` status in the VCF filter column. Indels and duplicated positions were removed; for multi-allelic variants, the most frequent alternate allele was retained. Variants with genotype missingness $> 5\%$ were excluded.

To enable cross-cohort projection, the reference panel was intersected with the genotyping array content of each target dataset. After intersection, variants with minor allele frequency (MAF) $< 1\%$ were removed and linkage disequilibrium (LD) pruning was performed using PLINK (`--indep-pairwise 500 50 0.5`). Low-complexity regions [41] and the HLA/MHC locus were excluded.

Samples were filtered from 4,150 to 4,094 individuals based on gnomAD metadata (downloaded January 1, 2025), excluding samples flagged as contaminated (n=2), hard-filtered (n=31), or identified as PCA outliers (n=23). A maximal subset of 3,400 unrelated individuals was selected using KING relatedness estimates.

##### S1.4.2 UK Biobank

All preprocessing and harmonization steps were implemented using a reproducible pipeline (available at `https://github.com/MattScicluna/manifold_genetics`).

UK Biobank genotypes were generated using the UK Biobank Axiom array. The dataset was then imputed using HRC reference panel and made available by the UK Biobank consortium and given the European bias of the Axiom array, the imputed dataset was selected to conduct the analysis. The imputed genomic coordinates were lifted to GRCh38 where required and were filtered for imputation quality (INFO $\geq 0.8$ and genotype posterior probability $\geq$ 0.9). Analyses were restricted to autosomal SNPs; indels and duplicated positions were removed, retaining the probe with lowest missingness where applicable. Variants with genotype missingness $> 5\%$ were excluded and a Hardy–Weinberg equilibrium filter was applied (`--hwe 1e-6 midp`).

Alleles were harmonized to the HGDP+1KGP reference using `bcftools +fixref`, and residual strand inconsistencies were removed using Will Rayner's pre-imputation harmonization pipeline with TOPMed allele frequencies as reference. SNPs were intersected with a version of the HGDP+1KGP panel filtered to $\leq 5\%$ missingness and MAF $\geq 0.01$.

An LD pruning step (`--indep-pairwise 150 kb 1 0.05`) was done as well as the removal of HLA/MHC and Genome In a Bottle low-complexity regions (`GRCh38_alldifficultregions.bed` file [42]).

For cross-cohort analyses, PCA and admixture models were trained on the HGDP+1KGP reference panel and applied to UK Biobank samples. For within-cohort visualization, a subset of 59,265 unrelated samples was used. Here, we subsampled to 10,000 White British and 5,000 Irish individuals, following [5]. PCs were computed using FlashPCA [43], admixture proportions were estimated for $K = 2$–10, and manifold embeddings were computed on PCA coordinates.

##### S1.4.3 All of Us

This study used data from the All of Us Research Program Controlled Tier Dataset v7, accessed through the All of Us Researcher Workbench.

Genotypes were restricted to autosomal SNPs, excluding indels and variants with genotype missingness $> 5\%$. Duplicate and multi-allelic positions were resolved as described for the reference panel. Alleles were harmonized to the HGDP+1KGP reference using `bcftools +fixref`, and strand inconsistencies were removed using Rayner's pre-imputation harmonization pipeline with TOPMed allele frequencies as reference.

As with UK Biobank, cross-cohort analyses projected All of Us samples onto PCA and admixture models trained on the HGDP+1KGP reference panel, whereas within-cohort visualization used a subsampled unrelated fit set of 61,711 individuals.

#### S1.5 Effect of neighborhood scale on diffusion embeddings

In this section we show how one can inadvertently produce distorted PHATE embeddings via misconstruction of the diffusion operator. Specifically, we show how extrinsic topological properties of the diffusion operator lead to distorted embeddings.

First we show that a disconnected diffusion operator produces embeddings that capture the disconnected components and their relative sizes at the expense of global structure. This can happen even if the underlying Markov process is irreducible, if the neighborhood size `KNN` is too small. Note that for our HGDP+1KGP dataset, the default settings of PHATE (`KNN` = 5) produce a disconnected diffusion operator, and we require `KNN` ≈ 100 to get an ergodic diffusion process. Then we show that connectivity of the diffusion operator is insufficient: if the operator is "too connected", then PHATE embeddings degenerate. We prove each of these below and demonstrate the effect on both simulated and real data.

##### S1.5.1 Case 1: Disconnected Diffusion Operator

We first show that between-component potential distances admit a lower bound that scales linearly with the component sizes. Then we show that, under mild boundedness assumptions and for sufficiently small $\epsilon$, distances between disconnected components are strictly larger than within-component distances.

Let $P$ be our $t$ step diffusion operator defined in Equation E1, suppressing the dependence on $t$ for notational clarity. Assume that our diffusion operator contains $> 1$ connected components. Recall that the PHATE implementation sets kernel values less than some threshold $p_{min}$ as 0 and adds a small value $\epsilon$ to all values of $P$. We assume $\epsilon < p_{min}$.

**Proposition 1** Let $x \in C_x$ and $y \in C_y$, where $C_x \neq C_y$ are internally fully-connected components (cliques) of $P$ that are disconnected from each other. Then

$$\mathcal{D}(x,y)^2 \geq (|C_x| + |C_y|) \log^2 \left( \frac{\epsilon}{p_{min} + \epsilon} \right).$$

*Proof* Since $x$ and $y$ belong to distinct disconnected components,

$$\mathcal{D}(x,y)^2 = \sum_{z \in C_x} \log^2 \left( \frac{\epsilon}{p_x(z) + \epsilon} \right) + \sum_{z \in C_y} \log^2 \left( \frac{\epsilon}{p_y(z) + \epsilon} \right).$$

We first bound $\log^2 \left( \frac{\epsilon}{p_x(z)+\epsilon} \right)$ and then use a symmetric argument for $\log^2 \left( \frac{\epsilon}{p_y(z)+\epsilon} \right)$. Combining $p_x(z) \geq p_{min}$ and $\epsilon < p_{min}$, we have

$$\frac{\epsilon}{p_x(z) + \epsilon} \leq \frac{\epsilon}{p_{min} + \epsilon} < 1 \tag{E8}$$

$$\Rightarrow \log^2 \left( \frac{\epsilon}{p_x(z) + \epsilon} \right) \geq \log^2 \left( \frac{\epsilon}{p_{min} + \epsilon} \right) \tag{E9}$$

$$\Rightarrow \sum_{z \in C_x} \log^2 \left( \frac{\epsilon}{p_x(z) + \epsilon} \right) \geq |C_x| \log^2 \left( \frac{\epsilon}{p_{min} + \epsilon} \right). \tag{E10}$$

Combining this with a symmetric argument for $|C_y|$ yields the desired conclusion. □

We note if $C_x, C_y$ are not fully connected, then we can still derive a bound

$$\mathcal{D}(x,y)^2 \geq (|S_x| + |S_y|) \log^2 \left( \frac{\epsilon}{p_{min} + \epsilon} \right),$$

where $S_x \subseteq C_x$ is the set of points where $p_x(z) > 0$ (and defined symmetrically for $S_y$). Finally, we note that $S_x \approx C_x$ in practice, since the $t$-step diffusion process will cause the intra-component points to become heavily connected.

**Proposition 2** *Global Separation of Components.* Let $x, y \in C_x$ and $y' \in C_y$ where $C_x, C_y$ are disconnected cliques. For a sufficiently small $\epsilon > 0$, the within-component potential distance is strictly smaller than the between-component potential distance:

$$\mathcal{D}(x, y) < \mathcal{D}(x, y').$$

*Proof* For points $x, y$ within the same clique $C_x$, the transition probabilities $p_x(z)$ and $p_y(z)$ are bounded between $p_{\min}$ and 1. The squared potential distance is thus bounded by the maximum internal contrast:

$$\mathcal{D}(x, y)^2 = \sum_{z \in C_x} \log^2 \left( \frac{p_y(z) + \epsilon}{p_x(z) + \epsilon} \right) \leq |C_x| \log^2 \left( \frac{1 + \epsilon}{p_{\min} + \epsilon} \right).$$

As $\epsilon \to 0$, this upper bound converges to the finite constant $|C_x| \log^2(1/p_{\min})$. In contrast, Proposition 1 establishes that the between-component distance $\mathcal{D}(x, y')^2$ is bounded below by a term proportional to $\log^2(\frac{\epsilon}{p_{\min}+\epsilon})$. As $\epsilon \to 0$, this lower bound approaches infinity:

$$\lim_{\epsilon \to 0} (|C_x| + |C_y|) \log^2 \left( \frac{\epsilon}{p_{\min} + \epsilon} \right) = \infty.$$

Since the within-component distance remains bounded while the between-component distance grows without bound as $\epsilon$ decreases, there exists a threshold $\epsilon^*$ such that for all $\epsilon < \epsilon^*$, we have $\mathcal{D}(x, y)^2 < \mathcal{D}(x, y')^2$. □

While $\epsilon$ is fixed in practice, this argument shows that for sufficiently small numerical regularization (relative to $p_{\min}$), between-component distances dominate within-component variation.

##### S1.5.2 Case 2: Overly-Connected Diffusion Operator

We now show that an overly connected diffusion operator yields PHATE embeddings devoid of meaningful information. First we show that increasing neighborhood connectivity causes the diffusion operator to become uniform. We then show that when the diffusion operator becomes uniform, PHATE embeddings collapse.

**Lemma 1** *Uniformization of the Diffusion Operator under Increasing Connectivity.* Let $A \in \mathbb{R}^{N \times N}$ be a nonnegative affinity matrix bounded above by $a_{\max}$. Assume $A_{x,y} \geq a_{\min} > 0$ if $y$ is among the $k_N$-nearest neighbors of $x$, and $A_{x,y} = 0$ otherwise. Let the diffusion operator $P$ be defined row-wise by $p_x(z) = A_{x,z}/d_x$, where $d_x = \sum_y A_{x,y}$. Then as $k_N \to N$ and $N \to \infty$,

$$\max_{x,z} \left| p_x(z) - \frac{1}{N} \right| \to 0$$

*Proof* For any $x$, the row sum $d_x$ is bounded below by the minimum affinity times the neighborhood size:

$$d_x = \sum_z A_{x,z} \geq a_{\min} k_N$$

Let $m_x = \min_z p_x(z)$ and $M_x = \max_z p_x(z)$. Since the probabilities in row $x$ sum to 1 over $N$ elements, the uniform probability $1/N$ must lie between the minimum and maximum:

$$m_x \leq \frac{1}{N} \leq M_x$$

Therefore, for any $z$, the absolute difference between $p_x(z)$ and the uniform distribution is bounded by the maximum spread of the probabilities:

$$\left| p_x(z) - \frac{1}{N} \right| \leq M_x - m_x \leq M_x$$

Because $p_x(z) = A_{x,z}/d_x$ and $A_{x,z} \leq a_{\max}$, we can uniformly bound $M_x$ for all $x$:

$$M_x \leq \frac{a_{\max}}{d_x} \leq \frac{a_{\max}}{a_{\min} k_N}$$

Combining these inequalities, we obtain a uniform bound across all $x$ and $z$:

$$\max_{x,z} \left| p_x(z) - \frac{1}{N} \right| \leq \frac{a_{\max}}{a_{\min} k_N}$$

As $k_N \to N$ and $N \to \infty$ (which implies $k_N \to \infty$), the term on the right approaches 0, completing the proof. □

For the PHATE $\alpha$-decaying kernel, if $k$ is large enough that $\|x - y\|_2 \leq \varepsilon_k(x)$ and $\|x - y\|_2 \leq \varepsilon_k(y)$ for all $x, y$, then

$$e^{-1} \leq k_{k,\alpha}(x, y) \leq 1$$

Thus Lemma 1 applies with $a_{\min} = e^{-1}$ and $a_{\max} = 1$.

**Proposition 3** *PHATE Collapse under Uniform Diffusion.* Suppose $\max_{x,z} \left| p_x(z) - \frac{1}{N} \right| \to 0$ as $k_N \to N$ and $N \to \infty$. Then the pairwise potential distances vanish ($\mathcal{D}(x, y)^2 \to 0$), and the PHATE embedding collapses to a single point.

*Proof* If the 1-step transition probabilities $p_x(z) \to \frac{1}{N}$, then the uniform distribution is stationary, meaning the $t$-step probabilities also converge uniformly: $p_x^t(z) \to \frac{1}{N}$ for all $x$ and $z$. Recall that the potential representation is defined element-wise as $U_x(z) = -\log(p_x^t(z))$. Substituting the limit, $U_x$ becomes a vector of identical constants, completely independent of the starting point $x$:

$$U_x \to \left( -\log\left(\frac{1}{N}\right), \ldots, -\log\left(\frac{1}{N}\right) \right)$$

Because $U_x$ converges to the exact same vector for all $x \in \mathcal{X}$, the potential distances $\mathcal{D}(x, y)^2 = \|U_x - U_y\|_2^2$ evaluate to zero. Applying MDS to a zero-distance matrix trivially yields a degenerate embedding where all points coincide. □

#### S1.6 Effect of population imbalance on embeddings

In the previous sections, we showed how intrinsic properties of the diffusion operator—namely, its connectivity and spectral collapse—can distort PHATE embeddings. We now turn to extrinsic dataset properties, specifically sampling design. We show that when population sizes are imbalanced, both PCA and PHATE produce embeddings that are distorted by the relative sample sizes. PCA distortion based on sample size has been well reported [11]. However, to the best of our knowledge, this has yet to be reported in PHATE.

We demonstrate this by showing that the core operators of both methods are explicit functions of the population counts, meaning the resulting embeddings capture sampling artifacts in addition to biological signal.

Let our dataset consist of $N$ samples drawn from $C$ distinct populations. Let $y_i \in \{1, \ldots, C\}$ denote the population label of sample $x_i$, and let $N_c = |\{i : y_i = c\}|$ be the sample size of population $c$. The relative proportion of each population is $\hat{\pi}_c = N_c/N$. Under the mixture model $x_i \mid y_i = c \sim p_c$, let:

$$\mu_c = \mathbb{E}[X \mid Y = c], \qquad \Sigma_c = \mathrm{Cov}(X \mid Y = c).$$

***PCA.***

Let the empirical mean and covariance matrix of the dataset be

$$\bar{x} = \frac{1}{N}\sum_{i=1}^{N} x_i, \qquad S = \frac{1}{N}\sum_{i=1}^{N}(x_i - \bar{x})(x_i - \bar{x})^\top.$$

Throughout this section, expectations are taken conditional on the observed labels $y$, so that the sampling proportions $\hat{\pi}_c = N_c/N$ are treated as fixed. The conditional expectation of the sample mean is:

$$\mathbb{E}[\bar{x} \mid y] := \mu_{\hat{\pi}} = \sum_{c=1}^{C} \hat{\pi}_c \mu_c.$$

By the law of total covariance applied to the mixture defined by $\hat{\pi}$, the conditional expectation of the sample covariance is

$$\mathbb{E}[S \mid y] = \sum_{c=1}^{C} \hat{\pi}_c \Sigma_c + \sum_{c=1}^{C} \hat{\pi}_c (\mu_c - \mu_{\hat{\pi}})(\mu_c - \mu_{\hat{\pi}})^\top.$$

This decomposition clearly shows that the covariance matrix is an explicit function of the sampling proportions $\hat{\pi}_c$. If a population is heavily oversampled, the leading principal components (the eigenvectors of $S$) will rotate to maximize variance within the oversampled group, distorting the global geometric relationships.

***PHATE.***
PHATE constructs its embedding based on a Markov transition matrix $P = D^{-1}A$, where $A$ is the sample-sample affinity matrix and $D$ is the diagonal degree matrix with $d_i = \sum_{j=1}^{N} A_{ij}$.

To see how population sizes affect this operator, we can group the sum for the degree $d_i$ by population. Let $\bar{A}_{i,c}$ be the arithmetic mean of the affinities between sample $i$ and all samples in population $c$:

$$\bar{A}_{i,c} = \frac{1}{N_c} \sum_{j:y_j=c} A_{ij}$$

We can then rewrite the degree $d_i$ as:

$$d_i = \sum_{c=1}^{C} \sum_{j:y_j=c} A_{ij} = \sum_{c=1}^{C} N_c \bar{A}_{i,c}$$

The one-step transition probability from sample $i$ to sample $j$ is therefore:

$$P_{ij} = \frac{A_{ij}}{d_i} = \frac{A_{ij}}{\sum_{c=1}^{C} N_c \bar{A}_{i,c}}$$

Furthermore, the total probability mass moving from sample $i$ into population $c'$ is:

$$\sum_{j:y_j=c'} P_{ij} = \frac{N_{c'} \bar{A}_{i,c'}}{\sum_{c=1}^{C} N_c \bar{A}_{i,c}}$$

Thus, the PHATE diffusion process is explicitly parameterized by population sample sizes. Even for constant $\bar{A}_{i,c}$, an artificially inflated $N_{c'}$ directly increases the transition mass into population $c'$. Since the final embedding is computed from the multi-step operator $P^t$ via potential distances, this initial sampling bias compounds over successive diffusion steps. Ultimately, this artificially compresses the potential distances towards overrepresented populations, systematically distorting the learned manifold geometry.

#### S1.7 Hyperparameter sensitivity analyses

The PHATE parameterization described in Supplementary Section S1.1 contains several hyperparameters that affect the apparent scale and continuity of the resulting ancestry coordinates. However, these parameters act at different stages of the algorithm and are therefore not interchangeable. The neighborhood parameter `KNN` determines the neighborhood graph on which diffusion is performed, diffusion time $t$ controls how far information propagates across this graph through $\mathbf{P}^t$, and the decay parameter $\alpha$ controls how sharply affinities decrease with distance within the adaptive kernel. In our experiments, $\gamma$ was fixed at 1 (except for the DLAtree, following convention from the original PHATE paper [15]), so the sensitivity analyses focus on `KNN`, $t$, and decay.

Supplementary Section S1.5.1 shows that disconnected neighborhood graphs can cause PHATE to emphasize these disconnected components at the expense of preserving global structure. This can happen when `KNN` is small. Increasing $t$ or lowering decay can broaden diffusion over an existing graph, but cannot create the edges needed to connect the graph. Conversely, excessively increasing `KNN` leads to the diffusion operator converging to uniform probabilities and the corresponding embeddings becoming over-smoothed. The relevant question is therefore whether there is an intermediate neighborhood-scale regime where ancestry relationships are neither fragmented nor over-smoothed.

We first evaluated this behavior in HGDP+1KGP by varying `KNN`, $t$, and decay (Extended Data Fig. 1 and Supplementary Figure S1). At small neighborhood sizes, the ancestry coordinates remain fragmented and retain the floral artifact observed under near-default settings in Figure 2. Although increasing $t$ or lowering decay produces embeddings that more clearly show the global structure, disconnected or weakly connected populations still appear as separated clusters in the middle of the graph. Only increasing `KNN` removes this artifact.

In our balanced-sampling UKBB and AoU, intermediate-to-large values of `KNN` and $t$ (e.g. `KNN`=500 and $t$=50-500) recover more continuous admixture trajectories, whereas smaller values under-resolve these relationships among less-sampled groups and overly large scales compress the embedding (Extended Data Figs. 2 and 3). Neighborhood-scale comparisons across PHATE, UMAP, and t-SNE further show that scale sensitivity is not unique to PHATE (Supplementary Figures S4 and S5). Increasing neighborhood size generally reduces local fragmentation, but the methods differ in how well they preserve continuous admixture structure after this adjustment.

Together, these analyses show that the improved PHATE ancestry coordinates arise from a consistent intermediate-scale regime associated with diffusion-operator connectivity, rather than from arbitrary hyperparameter choices.

#### S1.8 Sampling Strategies Comparison

In the main text, we construct ancestry coordinates on population-balanced subsets to mitigate the strong ancestry imbalance present in UKBB and AoU.

Our primary balanced sampling strategy follows the subset used for PCA computation in the UK Biobank analyses. Specifically, we retained 10,000 White British individuals and 5,000 White Irish individuals, following [5]. This reduces the dominance of the largest ancestry group while preserving substantial representation from a closely related population. For AoU, we similarly retained 10,000 individuals each from the "White", "Hispanic or Latino", and "Black or African American" groups.

In Supplementary Figure S8, we compare PHATE plots generated using this label-based subsampling against two label-free sampling alternatives: uniform random sampling and geometric sketching (`geosketch`) [44]. In both cases, we selected subsets containing 60,000 individuals, similar in size to the population-balanced subsets (59,265 for UKBB and 61,711 for AoU). Geometric sketching selects representative samples from a low-dimensional embedding with the goal of preserving the global geometry of the data manifold. For all sampling strategies, the same 50-dimensional PCA representation was used as input.

As shown in Supplementary Figure S8, uniform random sampling remains dominated by the largest ancestry groups, whereas both balanced sampling and geometric sketching recover substantially more continuous ancestry structure, in particular, admixture clines between non-dominant populations. This suggests that manifold-aware sampling strategies such as geometric sketching may provide a label-free alternative for constructing ancestry coordinates in strongly imbalanced cohorts.

#### S1.9 Synthetic Data Experiment

To systematically evaluate manifold learning algorithms, we construct a suite of synthetic datasets that capture a range of geometric and topological properties. Each dataset is designed to provide controlled ground-truth structure, enabling quantitative assessment of algorithmic performance under varying conditions such as noise, dimensionality, and connectivity. The following subsections describe the construction and properties of each synthetic dataset used in our experiments.

##### S1.9.1 DLA Tree

We generated synthetic datasets to evaluate the ability of embedding methods to recover continuous population structure under controlled conditions. We used a graph-conditioned variant of diffusion-limited aggregation (DLA) in which samples were generated along the branches of a predefined directed tree $\mathcal{G} = (\mathcal{V}, \mathcal{E})$. This framework allows explicit control over key features relevant to population genetics, including branching structure, uneven sampling, and missing populations (gaps). In this setting, data points are generated along predefined branches, with gaps introduced by omitting selected segments from the observed data while retaining the underlying ground-truth structure. Gaussian noise is added to all samples to mimic measurement variability. To test robustness to discontinuous sampling, selected branches were omitted from the observed data while retaining the full latent tree as ground truth. Ground-truth relationships were defined by geodesic distances along this latent tree, computed as shortest-path distances between samples on $\mathcal{G}$ before gap exclusion. This provides a controlled setting with known ground-truth structure for evaluating embedding performance.

Diffusion-Limited Aggregation (DLA) trees model branching developmental trajectories via sequential random walks. While standard DLA tree datasets rely on recursive random attachments to generate branching topologies [15], we introduce a graph-conditioned variant to afford precise macroscopic control over the latent structure.

In this formulation, the dataset strictly adheres to a user-defined directed tree $\mathcal{G} = (\mathcal{V}, \mathcal{E})$, where each edge $e \in \mathcal{E}$ represents a distinct branch containing $L_e$ sequentially generated samples. For a branch $e$, points are generated as $\mathbf{x}_k^{(e)} = \mathbf{x}_1^{(e)} + \sum_{t=2}^{k} \boldsymbol{\delta}_t^{(e)}$ in an ambient space $\mathbb{R}^d$.

To ensure that multiple branches emanating from a shared parent node remain structurally distinct and do not collapse into one another, random walks are constrained to orthogonal feature subspaces. If a node has $k$ outgoing edges, the $d$ dimensions are partitioned into $k$ roughly equal, disjoint subsets $\mathcal{S}_e$. The random displacement vector is restricted such that $[\boldsymbol{\delta}_t^{(e)}]_j \sim \mathcal{U}(-1, \rho)$ if $j \in \mathcal{S}_e$, and 0 otherwise. The spatial position of a parent node serves as the initialization $\mathbf{x}_1^{(e)}$ for all its outgoing branches.

To test algorithmic robustness to discontinuous sampling, we introduce missing populations (gaps) through topological exclusion. We completely omit a predefined subset of gap edges $\mathcal{E}_{\text{gap}} \subset \mathcal{E}$ from the final observation, although they are still generated in the background to construct the complete ground-truth coordinates. Finally, independent Gaussian noise $\boldsymbol{\varepsilon}_i \sim \mathcal{N}(\mathbf{0}, \sigma^2 \mathbf{I})$ is added to all retained points, yielding the observed dataset:

$$\mathcal{Y} = \left\{ \mathbf{x}_i^{(e)} + \boldsymbol{\varepsilon}_i \;\middle|\; e \in \mathcal{E} \setminus \mathcal{E}_{\text{gap}},\ i \in \{1, \ldots, L_e\} \right\}. \tag{E11}$$

Ground-truth geometry is defined by intrinsic continuous trajectories rather than ambient Euclidean distances. We construct a reference graph connecting consecutive points along all original branches (prior to gap exclusion), with edge weights given by unperturbed Euclidean distances $w_{ij} = \|\mathbf{x}_i - \mathbf{x}_j\|_2$. The ground-truth geodesic distance $d_{\text{geo}}(i, j)$ is defined as the shortest-path distance on this reference graph. Consequently, evaluating on this dataset tests an algorithm's capacity to infer latent connectivity across missing populations and recover the true intrinsic geometry.

##### S1.9.2 Swissroll

The synthetic Swissroll dataset represents a two-dimensional manifold $(y, t)$ smoothly embedded into a three-dimensional space $(x, y, z)$ via a spiral transformation. Multiple distributions are defined along the manifold, each with a variable number of points. For each distribution, random means for $t$ and $y$ are sampled, with Gaussian noise added to introduce local variability. These $(y, t)$ coordinates define the intrinsic geometry of the manifold. The 2D manifold is embedded into 3D space as follows:

$$x = t\cos(t), \quad y = y, \quad z = t\sin(t)$$

where the $y$ coordinate retains its noisy values. Following this transformation, additive Gaussian noise is applied to all three coordinates to simulate observation or measurement error.

To further challenge the robustness of manifold learning algorithms, we introduced a variant of the Swissroll containing significant discontinuities along the manifold. To generate these gaps, the means of the distributions are first sorted along the $t$ axis to ensure a consistent ordering. We then select $n_{\text{gaps}}$ indices and shift the $t$ means of all subsequent distributions by a fixed fraction of the total $t$ span. Mathematically, if the sorted $t$ means are denoted by $t_1, t_2, \ldots, t_N$, a gap of width $\Delta t$ inserted after index $i$ results in:

$$t_j \leftarrow t_j + \Delta t, \quad \forall j > i$$

This procedure creates regions along the spiral devoid of data points, simulating missing populations or fragmented manifolds.

Finally, to simulate high-dimensional real-world scenarios, the dataset is projected into a 100-dimensional space using a random orthogonal matrix. This transformation preserves pairwise distances and intrinsic geometry while eliminating axis alignments.

For both the standard and gap-augmented Swissrolls, ground-truth geodesic distances are computed in the unrolled $(u_t, y)$ coordinate space. The unrolling transformation is defined as:

$$u_t = 0.5 \left( \sqrt{t^2 + 1} \cdot t + \sinh^{-1}(t) \right)$$

Pairwise Euclidean distances in this $(u_t, y)$ space provide the intrinsic manifold distances used to evaluate algorithm performance.

##### S1.9.3 Toroidal Surface

The Toroidal Surface dataset simulates a toroidal (doughnut-shaped) structure, incorporating periodic boundary conditions to model the intrinsic geometry of the torus. Points are uniformly distributed across the toroidal surface, each initially defined by two angular coordinates, $\theta$ and $\phi$, sampled uniformly from the interval $[0, 2\pi)$. To introduce local variability, random noise is applied to these angular coordinates prior to spatial embedding.

The intrinsic $(\theta, \phi)$ coordinates are mapped into a three-dimensional Cartesian space using the standard torus parameterization:

$$\begin{aligned} x &= (R + r\cos\phi)\cos\theta \\ y &= (R + r\cos\phi)\sin\theta \\ z &= r\sin\phi \end{aligned}$$

where $R$ denotes the major radius and $r$ denotes the minor radius. To simulate measurement noise, additive Gaussian noise is applied to the resulting $(x, y, z)$ coordinates.

To evaluate the robustness of manifold learning algorithms against topological fragmentation, we introduce discontinuities by partitioning the toroidal surface into discrete angular bins. By selectively removing all samples within a subset of these bins, we generate a fragmented manifold characterized by sharp boundaries and disjoint regions. This procedure is designed to simulate data discontinuities and non-contiguous manifold structures, which pose significant challenges for methods that rely on local neighborhood preservation. Furthermore, to eliminate axis-alignment biases, the data is projected into a 100-dimensional space via a random orthogonal transformation.

Pairwise geodesic distances between points on the toroidal surface are computed in the angular coordinate space, taking periodic boundary conditions into account. The geodesic distance $d$ between two points $(\theta_1, \phi_1)$ and $(\theta_2, \phi_2)$ is defined as:

$$d = \sqrt{(R\Delta\theta)^2 + (r\Delta\phi)^2}$$

where $\Delta\theta$ and $\Delta\phi$ represent the minimal angular differences under periodic boundary conditions. This metric represents the shortest path along the surface, providing a basis for quantitative performance evaluation.

##### S1.9.4 Saddle Surface

The Saddle Surface dataset is modeled as a hyperbolic paraboloid. The dataset is constructed by sampling points from multiple Gaussian distributions distributed across the manifold, allowing for flexible specification of both the number of distributions and the density of points per distribution. Specifically, the dataset is generated by sampling mean values for parameters $u$ and $v$ uniformly within a defined bounded region. To introduce local variability, additive Gaussian noise is applied to both the $u$ and $v$ coordinates. The resulting latent $(u, v)$ pairs are mapped into a three-dimensional Cartesian space $(x, y, z)$ according to the following parameterization:

$$x = u, \quad y = v, \quad z = au^2 - bv^2$$

where $a$ and $b$ serve as curvature parameters. The Cartesian coordinates are further subjected to Gaussian noise.

To introduce topological discontinuities, gaps are generated by randomly selecting non-overlapping intervals along the $u$-axis and excluding any samples falling within these regions, resulting in a surface with distinct voids. Furthermore, the dataset is embedded into a 100-dimensional space via a random orthogonal transformation.

The intrinsic geometry of the saddle surface is benchmarked using pairwise geodesic distances. These are approximated by the Euclidean distance in the latent $(u, v)$ parameter space:

$$d = \sqrt{(u_1 - u_2)^2 + (v_1 - v_2)^2}$$

This metric serves as a ground-truth representation of the manifold's intrinsic structure, providing a basis for quantitative performance evaluation.

#### S1.10 PHATE Random Landmarking Benchmarking

We benchmark the default spectral landmark selection strategy in PHATE (PHATE v1) against our randomized landmarking approach (PHATE v2) in terms of computational efficiency and embedding fidelity. Specifically, we evaluate performance as a function of the neighborhood size `KNN`, the number of samples $n_{\text{sample}}$, and the number of landmarks $n_{\text{landmark}}$. Runtime is decomposed into the landmark operator and total execution time, while embedding quality is assessed by the sum of squared errors (SSE) between the exact PHATE embedding (computed without landmarking) and landmark-based embeddings after Procrustes alignment [45].

All experiments were conducted using 6 independent random seeds; reported runtime and quality metrics correspond to averages across runs.

##### S1.10.1 Scalability with Increasing Sample Size

Supplementary Figure S10a shows the runtime when varying the number of samples ($n_{\text{sample}} \in \{5{,}000, 10{,}000, 20{,}000, 50{,}000, 59{,}265\}$), evaluated on the UK Biobank balanced fitting subset (total $n = 59{,}265$). Unless otherwise specified, the `KNN` parameter was set to 10.

The randomized method scales much better than the default spectral approach: both total runtime and landmark operator time increase approximately linearly with $n_{\text{sample}}$ but with a markedly lower slope, remaining under 6 seconds at the full subset size, whereas the default method requires more than twice the runtime at the same scale.

##### S1.10.2 Runtime Analysis with Different KNN Values

Supplementary Figure S10b compares the runtime components of the default spectral approach and our randomized landmark selection across different KNN values ($k \in \{5, 25, 50, 100, 500, 1000, 2000, 4000\}$), evaluated on the UK Biobank balanced fitting subset ($n = 59{,}265$). The total runtime, as well as the time spent in the landmark operator, increases significantly with `KNN` for the default method, whereas the randomized version remains nearly constant across neighborhood sizes. This is expected since the randomized method avoids iterative SVD and KMeans computations, reducing the complexity from $\mathcal{O}(n^2k + nkt)$ to $\mathcal{O}(n \cdot n_{\text{landmark}})$. The speedup is visible in the "Landmark operator" components and total runtime, where the default method becomes over 3× slower for $k = 500$, confirming the efficiency of our approach for large KNN values.

##### S1.10.3 Embedding Fidelity with Increasing Number of Landmarks

Supplementary Figure S10c compares the sum of squared errors (SSE) between the exact PHATE embedding ($n_{\text{landmark}} = \text{None}$) and its versions using the landmark-building approach, for different numbers of landmarks ($n_{\text{landmark}}$ on the x-axis, SSE on the y-axis), evaluated on the UK Biobank balanced fitting subset. The SSE is computed using Procrustes analysis, following the procedure described in the original paper [15]. Both the default spectral method and our randomized version show a decreasing SSE as $n_{\text{landmark}}$ increases, indicating a better approximation with more landmarks. Notably, the randomized approach achieves comparable or slightly lower SSE than the default method for most values of $n_{\text{landmark}}$, confirming that the gain in computational efficiency does not come at the expense of embedding quality (the overall curve closely matches the one reported in [15]).

##### S1.10.4 Comparative scaling with sample size and neighborhood scale across methods

Supplementary Figure S10d–e compare total runtime scaling across PHATE v1 (spectral landmarking), PHATE v2 (randomized landmarking), t-SNE, and UMAP on the UK Biobank balanced fitting subset. Panel d evaluates scaling with respect to the number of samples, while panel e evaluates scaling with respect to neighborhood size.

As the number of samples increases, PHATE v2 consistently exhibits the lowest total runtime among all methods, with a substantially flatter scaling curve compared to PHATE v1. In contrast, PHATE v1 shows noticeably steeper growth, and t-SNE exhibits the highest computational cost across sample sizes. UMAP scales more favorably than t-SNE but remains slower than PHATE v2. These results demonstrate that randomized landmarking substantially improves the scalability of PHATE with respect to dataset size.

A similar pattern is observed when increasing neighborhood size. PHATE v2 maintains the lowest runtime across the full range of neighborhood parameters, whereas PHATE v1 and UMAP show moderate growth and t-SNE scales most steeply. Notably, the computational gap widens at larger neighborhood sizes, highlighting the advantage of randomized landmarking in regimes that emphasize global structure.

Together, these results indicate that PHATE with randomized landmarking provides the most favorable scaling behavior across both sample size and neighborhood scale, supporting its applicability to large-scale biobank datasets.

##### S1.10.5 Scalability benchmarking on synthetic DLA data

Supplementary Figure S11a–b decompose runtime across KNN search, landmark operator, metric MDS, and remaining steps as functions of sample size and neighborhood size, respectively. As in the UK Biobank experiments, the landmark operator is the dominant computational bottleneck in PHATE v1, particularly at large neighborhood scales. Randomized landmarking substantially reduces this cost, leading to pronounced runtime improvements that become more evident as both the number of samples and neighborhood size increase. The relative contribution of KNN search and metric MDS remains similar across implementations, indicating that the primary gain arises from replacing the spectral landmark selection step.

Supplementary Figure S11c–d compare total runtime scaling across methods. PHATE v2 (Randomized landmarking) consistently achieves the lowest runtime across both increasing sample size and neighborhood scale, while PHATE v1 (spectral landmarking) exhibits steeper growth. UMAP shows the most pronounced increase with neighborhood scale, and t-SNE scales more moderately but remains slower than PHATE v2 at larger parameter values. Importantly, the magnitude and slope of these scaling curves differ from those observed in the UK Biobank subset.

These differences highlight that data topology substantially influences benchmarking outcomes. The UK Biobank subset reflects complex, high-density population structure with heterogeneous sampling, whereas the synthetic DLA dataset represents a controlled branched manifold with lower intrinsic dimensionality and simpler geometry. Such structural differences affect neighborhood graph connectivity, diffusion behavior, and the cost of spectral operations, thereby altering absolute runtimes and scaling slopes. Nevertheless, across both datasets, randomized landmarking consistently alleviates the dominant spectral bottleneck and preserves favorable scaling properties, demonstrating robustness to variation in underlying data topology.

### S2 Supplementary Figures

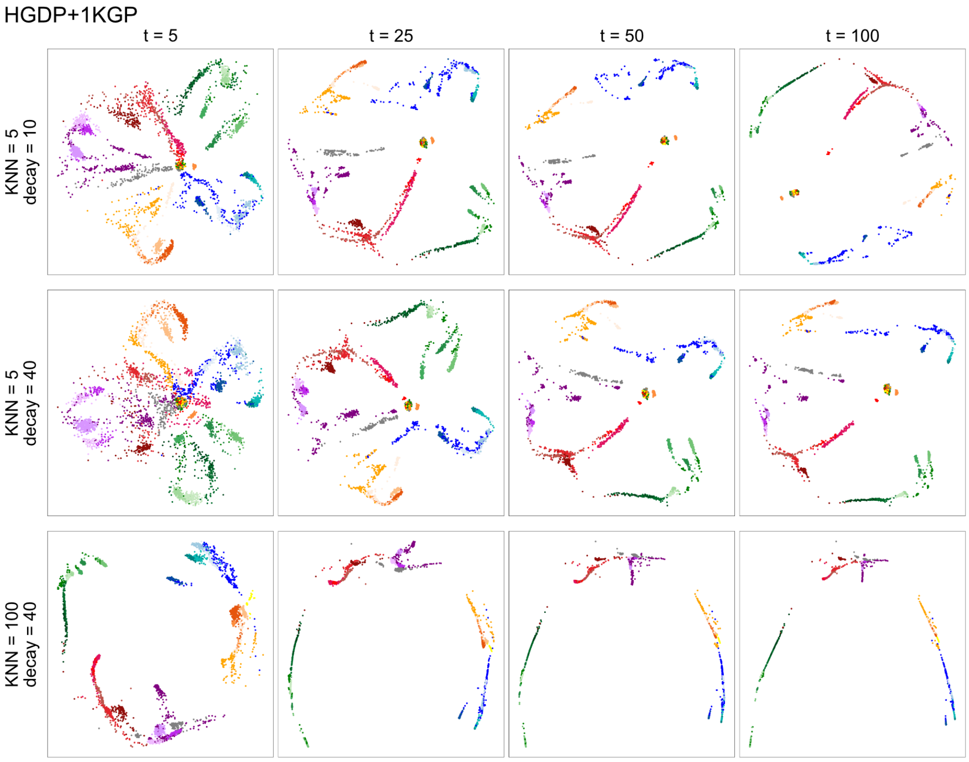


**Fig. S1** PHATE fitted to HGDP+1KGP ($N$ = 4,094) varying `KNN`, $t$, and decay. All other parameters are set to default.

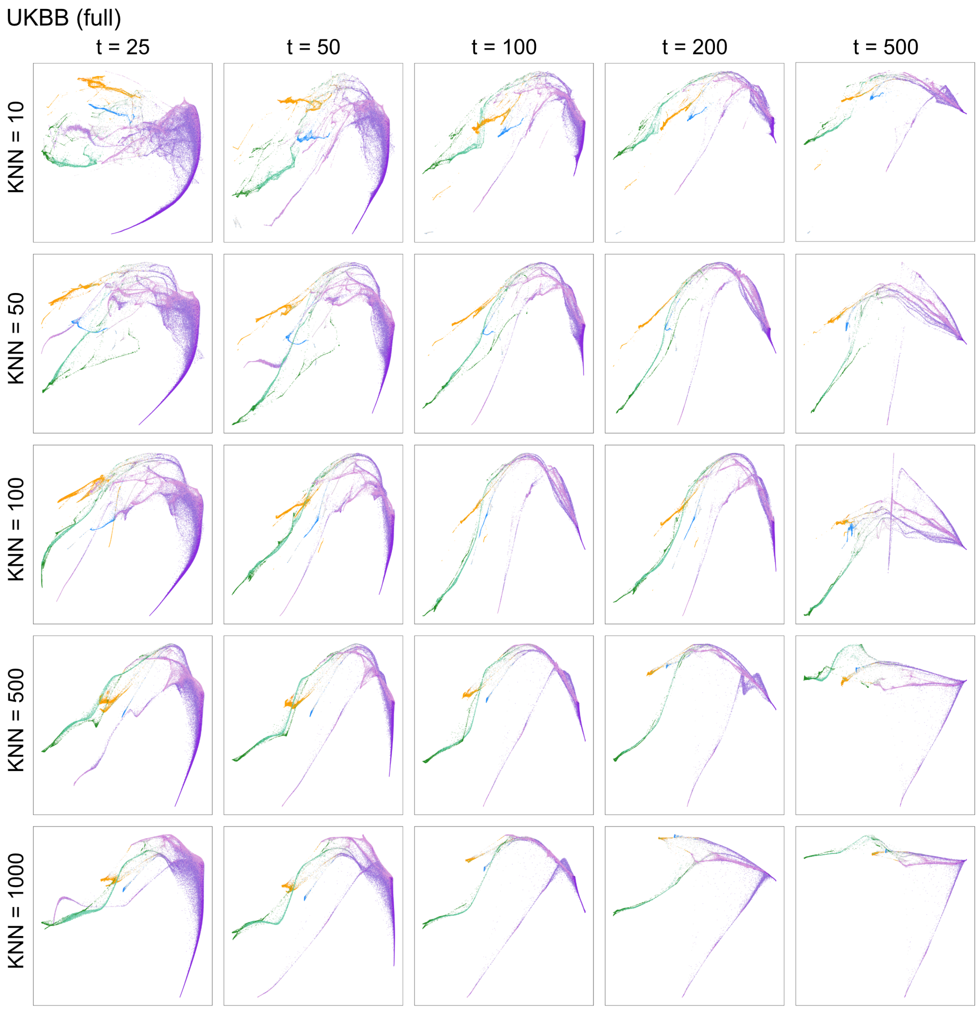


**Fig. S2** PHATE fitted to UKBB ($N$ = 486,748) varying `KNN` and $t$. All other parameters are set to default.

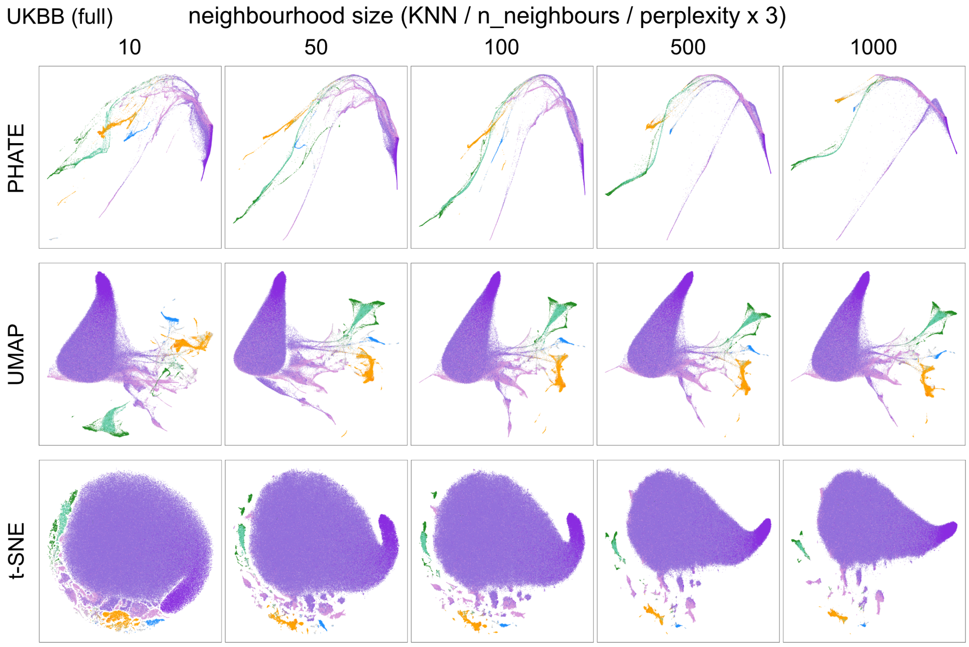


**Fig. S3** PHATE, UMAP, and t-SNE coordinates for UKBB ($N$ = 486,748) with increasing neighborhood size. PHATE was run with $t = 100$ and UMAP with `min_dist` = 0.5. All other parameters are set to default.

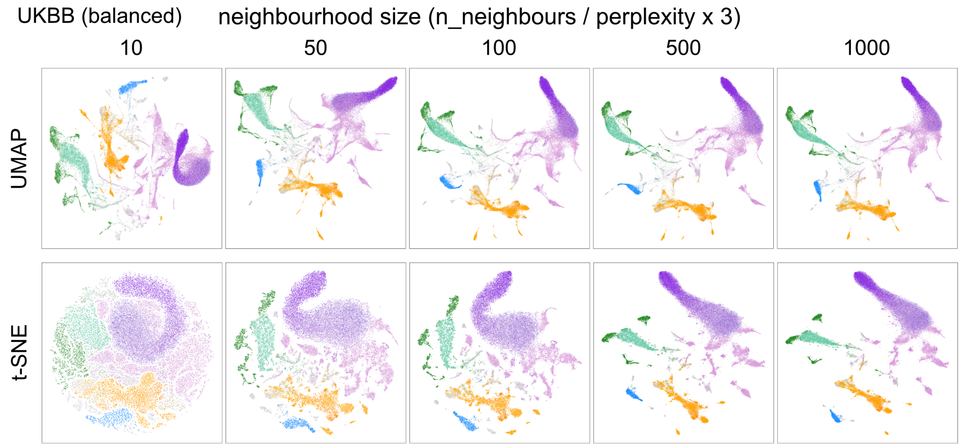


**Fig. S4** UMAP and t-SNE coordinates for balanced UKBB ($N = 59{,}265$) with increasing neighborhood size. UMAP was run with `min_dist` $= 0.5$. All other parameters are set to default.

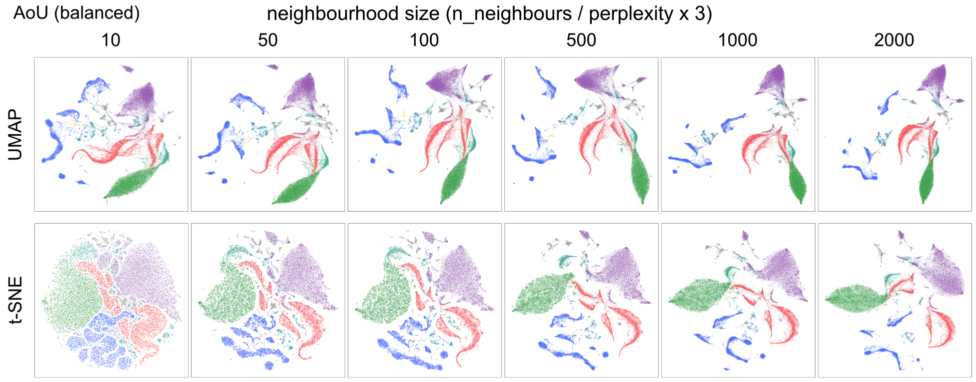


**Fig. S5** UMAP and t-SNE coordinates for balanced AoU ($N = 61{,}711$) with increasing neighborhood size. UMAP was run with `min_dist` $= 0.5$. All other parameters are set to default.

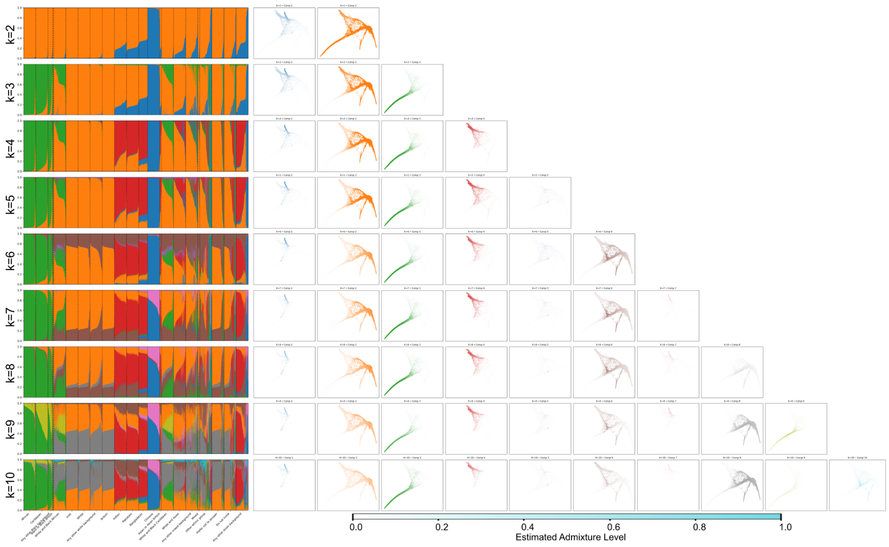


**Fig. S6** Balanced UKBB ($N$ = 59,265) admixture bar and PHATE plots colored by estimated admixture level. PHATE was run with `KNN`= 500 and $t = 100$. All other parameters are set to default.

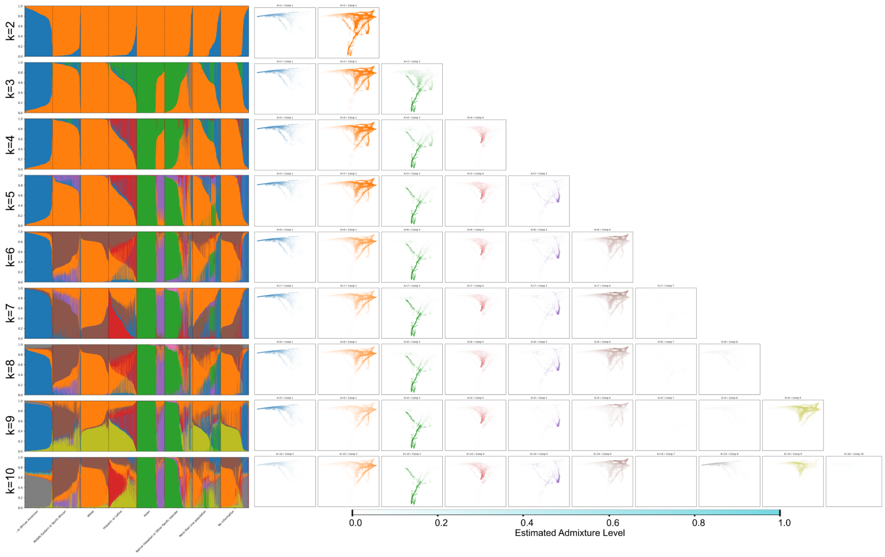


**Fig. S7** Balanced AoU ($N = 61{,}711$) admixture bar and PHATE plots colored by estimated admixture level. PHATE was run with `KNN`= 500 and $t = 100$. All other parameters are set to default.

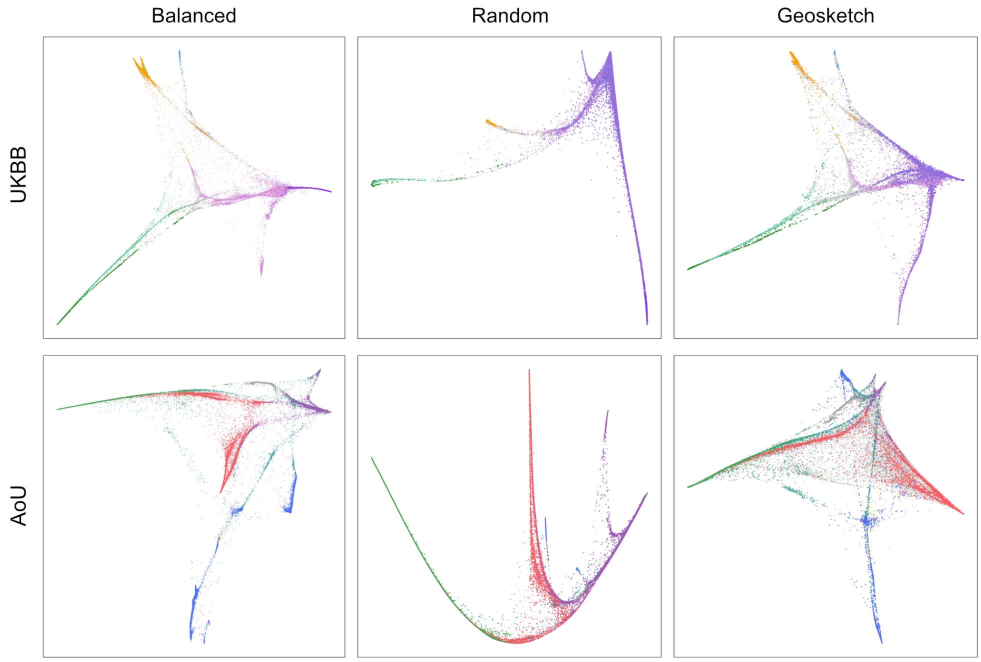


**Fig. S8** PHATE embeddings obtained after balanced sampling (UKBB $N$ = 59,265, AoU $N$ = 61,711), uniform random sampling ($N$ = 60,000), and geometric sketching ($N$ = 60,000). All embeddings were generated with PHATE using `KNN` = 500 and $t$ = 100. All other parameters are set to default.

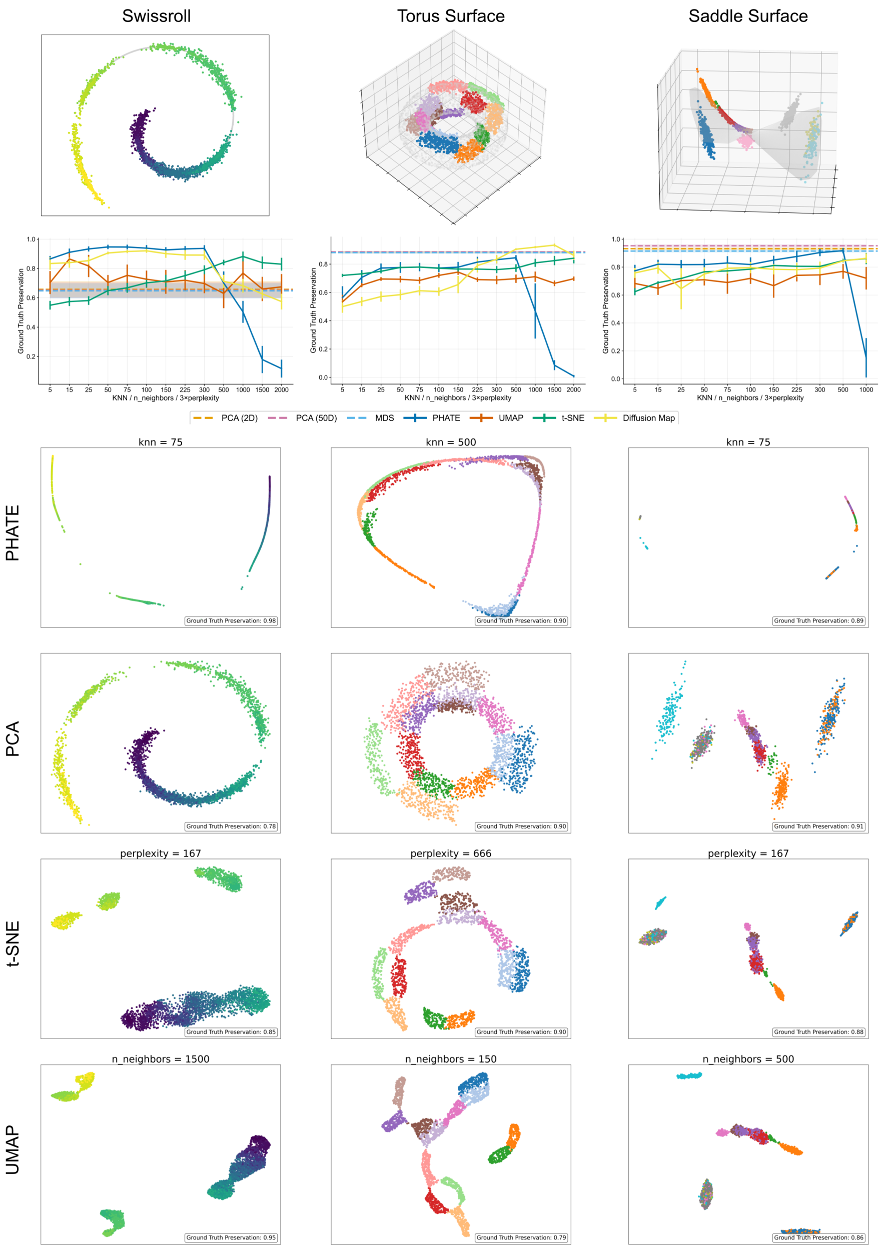


**Fig. S9** Additional synthetic benchmarks using distributions sampled on Swiss roll, toroidal surface, and saddle surface manifolds with gaps. For each dataset, we show the ground-truth geometry, ground-truth preservation across neighborhood scales, and representative embeddings from PHATE, PCA, t-SNE, and UMAP. These examples show that distortions under missing data and inappropriate neighborhood scale are not specific to the DLA tree, and support the need for careful scale selection when recovering continuous structure.

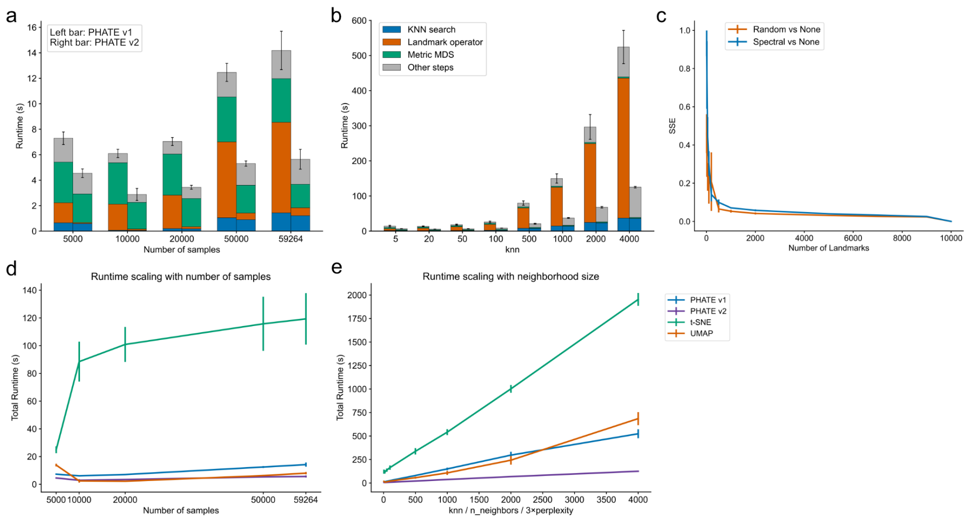


**Fig. S10** Randomized landmarking improves PHATE scalability while preserving embedding fidelity on the UKBB balanced fitting subset. Panels a and b share the same legend and runtime component color scheme. a) Runtime decomposition across KNN search, landmark operator, metric MDS, and remaining steps for the default spectral landmark selection (PHATE v1) and the randomized landmarking approach (PHATE v2) as a function of sample size. b) Runtime decomposition across the same components as a function of neighborhood size (KNN). The landmark operator is the dominant computational bottleneck in PHATE v1, whereas randomized landmarking substantially reduces this cost, resulting in increasing speedups with larger sample sizes and neighborhood scales. c) Embedding fidelity measured by SSE between the exact PHATE embedding (no landmarking) and landmark-based embeddings across landmark numbers. Randomized landmarking achieves embedding accuracy comparable to the default spectral method across landmark sizes. d) Total runtime scaling with increasing sample size for PHATE v1, PHATE v2, t-SNE, and UMAP. e) Total runtime scaling with neighborhood size across methods. PHATE v2 with randomized landmarking exhibits more favorable scaling with both sample size and neighborhood size than PHATE v1 and competing approaches.

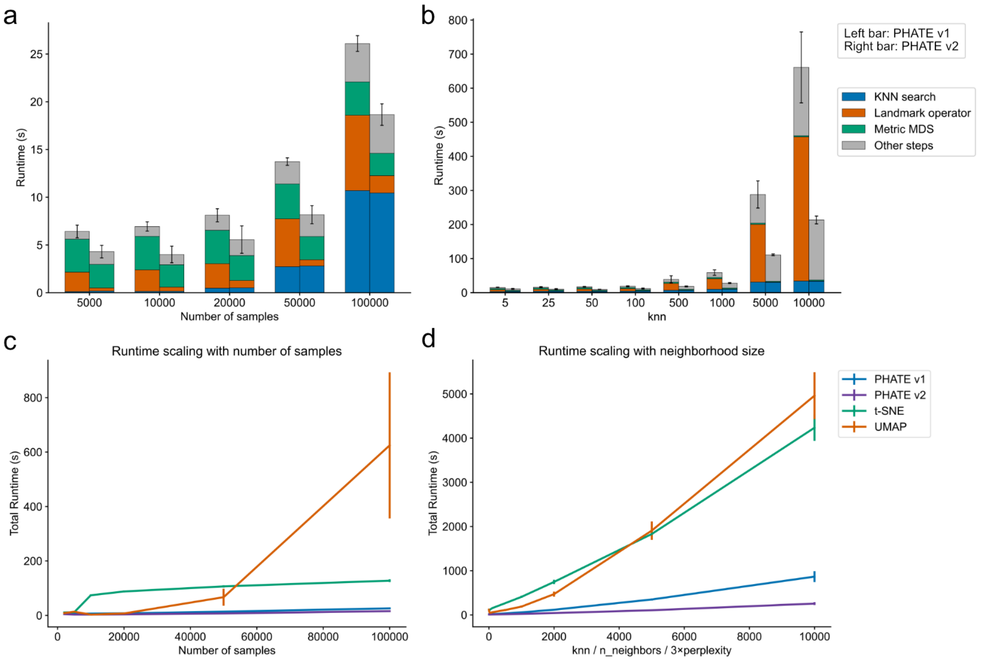


**Fig. S11** Randomized landmarking improves PHATE computational scalability on synthetic DLA data. a) Runtime decomposition across KNN search, landmark operator, metric MDS, and remaining steps for the default spectral landmark selection (PHATE v1) and the randomized landmarking approach (PHATE v2) as a function of sample size. b) Runtime decomposition across the same components as a function of neighborhood size. The landmark operator dominates the default implementation, whereas randomized landmarking substantially reduces this cost, leading to consistent speedups as both sample size and neighborhood size increase. c) Total runtime scaling with increasing number of samples for PHATE v1, PHATE v2, t-SNE, and UMAP. d) Total runtime scaling with increasing neighborhood size across methods. PHATE with randomized landmarking maintains the lowest runtime and exhibits the most favorable scaling behavior across both sample size and neighborhood scale.

### S3 Supplementary Tables

**Table S1** Populations included in and excluded from geographic preservation analysis in Section 2.3. The Geographic Preservation Metric is defined in Section 4.3.

| | |
|---|---|
| Used for Geographic Preservation Metric | Bantu Kenya, Bantu South Africa, Biaka Pygmy, Esan in Nigeria (ESN), Gambian in W. Gambia (GWD), Luhya in Webuye (LWK), Mende in Sierra Leone (MSL), Mandenka, Mbuti Pygmy, San, Yoruba in Ibadan (YRI), Yoruba (HGDP), Bengali in Bangladesh (BEB), Balochi, Brahui, Burusho, Gujarati Indian in Houston (GIH), Hazara, Indian Telugu in UK (ITU), Kalash, Makrani, Pathan, Punjabi in Lahore (PJL), Sri Lankan Tamil in UK (STU), Sindhi, Chinese Dai in Xishuangbanna (CDX), Han Chinese in Beijing (CHB), Southern Han Chinese (CHS), Han (HGDP), Japanese in Tokyo (JPT), Japanese (HGDP), Kinh Vietnamese in Ho Chi Minh City (KHV), Yakut, Adygei, Basque, Finnish (FIN), French (HGDP), British in England & Scotland (GBR), Iberian in Spain (IBS), Italian (HGDP), Orcadian, Russian (HGDP), Sardinian, Toscani in Italy (TSI), Tuscan (HGDP), Bedouin, Druze, Mozabite, Palestinian, Melanesian, Papuan |
| Excluded | African Caribbean in Barbados (ACB), African Ancestry in SW USA (ASW), Colombian in Medellín (CLM), Colombian (HGDP), Karitiana, Mexican Ancestry in LA (MXL), Maya (HGDP), Peruvian in Lima (PEL), Puerto Rican in Puerto Rico (PUR), Pima (HGDP), Surui (HGDP), Utah Residents (CEPH, CEU) |

**Table S2** Admixture Preservation Results

| | HGDP+1KGP | | | UKBB (balanced) | | | AoU (balanced) | | |
|---|---|---|---|---|---|---|---|---|---|
| KNN | PHATE | UMAP | t-SNE | PHATE | UMAP | t-SNE | PHATE | UMAP | t-SNE |
| 5 | $0.494 \pm 0.002$ | $0.375 \pm 0.023$ | $0.366 \pm 0.015$ | — | — | — | — | — | — |
| 10 | $0.554 \pm 0.002$ | $0.415 \pm 0.076$ | $0.371 \pm 0.006$ | $0.751 \pm 0.003$ | $0.669 \pm 0.030$ | $0.482 \pm 0.010$ | 0.745 | $0.716 \pm 0.009$ | $0.501 \pm 0.002$ |
| 20 | $0.604 \pm 0.001$ | $0.504 \pm 0.047$ | $0.359 \pm 0.023$ | $0.751 \pm 0.003$ | $0.693 \pm 0.009$ | $0.505 \pm 0.010$ | 0.753 | $0.718 \pm 0.014$ | $0.522 \pm 0.002$ |
| 50 | $0.672 \pm 0.001$ | $0.485 \pm 0.112$ | $0.396 \pm 0.008$ | $0.758 \pm 0.003$ | $0.721 \pm 0.010$ | $0.529 \pm 0.005$ | 0.786 | $0.721 \pm 0.013$ | $0.537 \pm 0.001$ |
| 100 | $0.717 \pm 0.001$ | $0.576 \pm 0.120$ | $0.424 \pm 0.003$ | $0.765 \pm 0.002$ | $0.729 \pm 0.014$ | $0.540 \pm 0.013$ | 0.794 | $0.736 \pm 0.012$ | $0.554 \pm 0.001$ |
| 500 | $0.750 \pm 0.001$ | $0.533 \pm 0.110$ | $0.540 \pm 0.002$ | $0.796 \pm 0.002$ | $0.734 \pm 0.007$ | $0.577 \pm 0.005$ | 0.820 | $0.762 \pm 0.023$ | $0.587 \pm 0.001$ |
| 1000 | $0.727 \pm 0.001$ | $0.596 \pm 0.063$ | $0.675 \pm 0.005$ | $0.793 \pm 0.002$ | $0.722 \pm 0.022$ | $0.636 \pm 0.003$ | 0.841 | $0.758 \pm 0.027$ | $0.601 \pm 0.001$ |
| 2000 | — | — | — | $0.855 \pm 0.000$ | $0.725 \pm 0.031$ | $0.665 \pm 0.016$ | 0.855 | $0.742 \pm 0.033$ | $0.618 \pm 0.004$ |
| 4000 | $-0.056 \pm 0.001$ | $0.038 \pm 0.007$ | $0.673 \pm 0.037$ | — | — | — | — | — | — |

**Table S3** HGDP+1KGP Geographic Preservation Results

| | HGDP+1KGP | | |
|---|---|---|---|
| KNN | PHATE | UMAP | t-SNE |
| 5 | $0.535 \pm 0.003$ | $0.479 \pm 0.057$ | $0.481 \pm 0.017$ |
| 10 | $0.591 \pm 0.002$ | $0.464 \pm 0.157$ | $0.471 \pm 0.014$ |
| 20 | $0.633 \pm 0.002$ | $0.443 \pm 0.148$ | $0.458 \pm 0.016$ |
| 50 | $0.684 \pm 0.003$ | $0.589 \pm 0.104$ | $0.496 \pm 0.021$ |
| 100 | $0.743 \pm 0.002$ | $0.618 \pm 0.194$ | $0.530 \pm 0.007$ |
| 500 | $0.764 \pm 0.003$ | $0.545 \pm 0.166$ | $0.699 \pm 0.004$ |
| 1000 | $0.769 \pm 0.002$ | $0.748 \pm 0.053$ | $0.700 \pm 0.003$ |
| 4000 | $-0.048 \pm 0.001$ | $0.019 \pm 0.002$ | $0.792 \pm 0.028$ |

**Table S4** UKBB (Full) Admixture Preservation Results

| | UKBB (Full) | | |
|---|---|---|---|
| KNN | PHATE | UMAP | t-SNE |
| 10 | 0.555 | 0.582 | 0.400 |
| 20 | 0.571 | 0.608 | 0.407 |
| 50 | 0.596 | 0.607 | 0.408 |
| 100 | 0.605 | 0.568 | 0.412 |
| 500 | 0.602 | 0.565 | 0.456 |
| 1000 | 0.575 | 0.564 | 0.490 |

[40] Damrich, S. & Hamprecht, F. Ranzato, M., Beygelzimer, A., Dauphin, Y., Liang, P. & Vaughan, J. W. (eds) *On umap's true loss function.* (eds Ranzato, M., Beygelzimer, A., Dauphin, Y., Liang, P. & Vaughan, J. W.) *Advances*

*in Neural Information Processing Systems*, Vol. 34, 5798–5809 (Curran Associates, Inc., 2021). URL https://proceedings.neurips.cc/paper_files/paper/2021/file/2de5d16682c3c35007e4e92982f1a2ba-Paper.pdf.
